# A Novel Stress Response Pathway Mediates Antibiotic Tolerance and Architecture in *Pseudomonas aeruginosa* Biofilms

**DOI:** 10.64898/2025.12.19.695352

**Authors:** Ainelen Piazza, Catriona M. A. Thompson, Govind Chandra, Harshanie Dasanayaka, Gerhard Saalbach, Carlo Martins, Eleftheria Trampari, Mark A. Webber, Freya Harrison, Jacob G. Malone

**Affiliations:** Department of Molecular Microbiology, John Innes Centre, Norwich, Norwich Research Park, NR4 7UH UK; The Centre for Microbial Interactions, Norwich, Norwich Research Park, NR4 7UG, UK; John Innes Centre Proteomics Platform; Quadram Institute Bioscience, Norwich Research Park, Norwich, Norfolk NR4 7UQ, UK; School of Life Sciences, Gibbet Hill Campus, The University of Warwick, Coventry, CV4 7AL, UK; School of Biological Sciences, University of East Anglia, Norwich Research Park, Norwich, NR4 7TJ, UK

## Abstract

*Pseudomonas aeruginosa* is a multidrug-resistant opportunistic pathogen, with chronic infections often associated with biofilm formation. Here, we investigate the previously uncharacterized gene *PA3049*, which is upregulated under biofilm conditions, to determine its role in infection, biofilm formation, and antimicrobial tolerance. We show that the small uncharacterised protein PA3049, renamed as <u>B</u>iofilm <u>a</u>ntibiotic tolerance <u>R</u>egulator (BatR), promotes biofilm establishment and enhances bacterial survival in sub-inhibitory concentrations of antibiotics. Proteomic analysis revealed that BatR influences the R2/F2 pyocin cluster, which drives explosive cell lysis and extracellular DNA (eDNA) release during biofilm development. We further identify a specific interaction between BatR and PA0486 (SrkA), an uncharacterised Ser/Thr protein kinase. We show that SrkA controls biofilm and pyocyanin production, and lysis-mediated eDNA release through regulation of the R2/F2 pyocin cluster and activation of bacteriophage Pf1. Our findings support a model in which SrkA directly regulates key biofilm-associated phenotypes, while BatR acts as a modulatory partner that tunes SrkA activity under specific conditions. Finally, BatR function was tested in high-validity infection models, including the *ex vivo* pig lung model of cystic fibrosis infection and a synthetic chronic-wound model. In these models, BatR contributes to biofilm architecture and antibiotic tolerance, and modulates pyocyanin production. Our study implicates the BatR/SrkA system in the response of *P. aeruginosa* biofilms to antibiotic challenge in lung infections.

## Introduction

*Pseudomonas aeruginosa* is a virulent, opportunistic human pathogen, designated as one of the ESKAPE group of bacteria (2) that are leading causes of multidrug-resistant, nosocomial bacterial infections. *P. aeruginosa* presents a significant challenge in healthcare settings where it causes highly persistent, chronic infections, primarily affecting catheter and cannula implant (3), burn or other major injury victims (4), chronic wounds (5), and immunocompromised patients (6). *P. aeruginosa* is also a major respiratory pathogen, causing both acute and chronic lung infections. Pulmonary infections caused by *P. aeruginosa* are a major cause of mortality and morbidity in people with the genetic disorder cystic fibrosis (CF) (7, 8).

Chronic *P. aeruginosa* infections are frequently associated with biofilms, which enable it to evade host immune responses and confer broad resistance/tolerance to antimicrobial agents, complicating treatment strategies. Bacterial biofilms exhibit common traits and phenotypic characteristics, including cell-to-cell communication (quorum sensing), the production and deployment of extracellular polymeric substances and extracellular DNA (eDNA), and the spatially structured control of motility, adhesins and cyclic di-GMP (c-di-GMP) levels (9). The upregulation of genes linked to stationary phase adaptation, environmental stress and anaerobiosis further underscores the distinct features associated with biofilm growth (10, 11). Additionally, the spatial arrangement of cells within the biofilm community, exposed to multiple resource gradients, introduces heterogeneity in cell physiology and metabolism that plays an important role in antibiotic tolerance. This diversity includes significant subpopulations of less metabolically active (dormant) cells, which contribute to the notable tolerance of biofilms to antibiotics designed to target active metabolic processes (12). For example, dormant cells within oxygen-depleted zones of *P. aeruginosa* biofilms exhibit lower overall mRNA transcript abundance and increased tolerance to ciprofloxacin and tobramycin (12).

A crucial aspect of biofilm-associated infections is the frequent exposure to sub-inhibitory concentrations (SICs) of antibiotics, which do not kill bacteria but can still influence community behaviour (13, 14). SICs often occur in tissues where antibiotic penetration is incomplete (15), in biofilms where the matrix limits drug diffusion (16), or when antibiotics are administered in lower doses to mitigate toxicity (17). SICs are particularly important in the context of biofilm-related infections because they can modulate bacterial gene expression, leading to the induction of stress responses (18), virulence (19), quorum sensing (19, 20) and further biofilm formation (21, 22).

eDNA is the most abundant polymer in the *P. aeruginosa* biofilm matrix and its production levels vary across strains. It plays a critical role by mediating cell-cell and cell-matrix interactions, thereby stabilising the biofilm architecture (23). There is no universal mechanism for eDNA production across different species. However, in the majority of cases eDNA is derived from genomic DNA released into the extracellular milieu as a consequence of cell death, a process that promotes biofilm formation by supporting the growth and adhesion of the surviving community (23). During the early stages of *P. aeruginosa* biofilm formation, eDNA is predominantly driven by explosive cell lysis, a programmed cell death mechanism mediated by the Lys endolysin, encoded within the R-pyocin and F-pyocin gene clusters (24). Pyocins are bacteriocins produced by *P. aeruginosa*, which contribute to interbacterial competition and survival within polymicrobial environments, but their associated lytic components also facilitate eDNA release (1, 24). As the biofilm matures, additional regulatory systems come into play. Quorum sensing (QS) and the activity of the filamentous Pf4 phage both trigger lysis events in a subpopulation of cells, maintaining eDNA production during later biofilm stages (25, 26). Other mechanisms also cause cell lysis and subsequent eDNA release via flagella and type IV pili (26). Furthermore, eDNA release in *P. aeruginosa* also occurs through oxidative stress caused by hydrogen peroxide (H_2_O_2_) generation mediated by pyocyanin production (27). Pyocyanin is a redox-reactive phenazine molecule that is produced by 90-95% of *P. aeruginosa* strains and is present in high concentrations in CF lung infections (28). These coordinated cell death mechanisms ensure continuous structural reinforcement and adaptation of the biofilm community.

Although a consensus has emerged on the role of many biofilm-associated traits, the large number of uncharacterized genes reported as differentially expressed under biofilm conditions highlights the substantial gaps that remain in our understanding of this complex bacterial lifestyle (29). Among the minority of upregulated loci in dormant, biofilm-dwelling *P. aeruginosa* cells is the small, uncharacterised gene *PA3049* (12), annotated as a homolog of the *Escherichia coli* <u>r</u>ibosome <u>m</u>odulator factor (*RMF*) (12). RMF, a ribosomally associated protein, facilitates ribosome hibernation by associating with 100S ribosome dimers and modulates *E. coli* translation during the stationary phase (30). However, while RMF is crucial for ribosome hibernation in *E. coli* (31), previous studies have demonstrated that PA3049 does not fulfil this function in *P. aeruginosa* (32). Despite its conservation in sequenced *P. aeruginosa* strains and presence in biofilm cells, the role of *PA3049* in *P. aeruginosa* biofilm formation remains unknown (32).

In this study, we characterise PA3049 and define its contribution to biofilm formation and antibiotic tolerance in *P. aeruginosa*. Phylogenetic analysis showed that *PA3049*-like genes are widespread among γ-proteobacteria, with a substantial degree of sequence and structural divergence predicted between *Pseudomonadales* and *Enterobacterales*. We show that PA3049 plays an important role in shaping biofilm architecture and enabling established biofilms to withstand sub-inhibitory antimicrobial challenge. Proteomic and phenotypic analyses linked PA3049 activity to pyocyanin and R2-F2 pyocin production, both systems involved in biofilm development. PA3049 interacts directly with the uncharacterised Ser/Thr kinase PA0486 (SrkA), which also modulates pyocyanin levels, biofilm formation, and cell lysis, suggesting an *in vivo* regulatory connection.

Finally, the clinical relevance of PA3049 function was demonstrated using two clinically validated infection models: an established *ex vivo* pig lung (EVPL) model for *P. aeruginosa* biofilm infection of CF bronchioles (33), and the synthetic chronic wounds model (SCW) for *P. aeruginosa* diabetic foot infections (34).

Taken together, our findings indicate that PA3049 has diverged from the ribosome-associated proteins found in Enterobacteria and functions as a specialised, posttranslational regulator of biofilm formation and stress adaptation in *P. aeruginosa*. Given its role in promoting biofilm-associated antibiotic tolerance, we renamed PA3049 as BatR (<u>B</u>iofilm <u>a</u>ntibiotic tolerance <u>R</u>egulator).

## Results

### *PA3049* homologs are widespread in γ-proteobacteria

To uncover the role of PA3049 (BatR) in *P. aeruginosa*, we first analysed the distribution of *batR*-like genes in bacterial genomes, and compared its predicted structure to *E. coli* RMF, which shares ∼49.09% sequence similarity with PA3049. Alignment of *batR* homologs from different *Pseudomonas* species with the *RMF* sequence from *E. coli* shows a 15-residue C-terminal extension encoded by *batR* that is absent from *RMF* (Fig 1a). AlphaFold3 (35) three-dimensional protein structure prediction of *P. aeruginosa* BatR highlighted a marked difference in the C-terminal fold compared to its *E. coli* homolog. Specifically, BatR contains a predicted 12 residue α-helix at the C terminus (red box), in addition to the two α helices connected by a 13-amino acid linker region that characterise *E. coli* RMF (Fig 1b).

**Fig 1.**
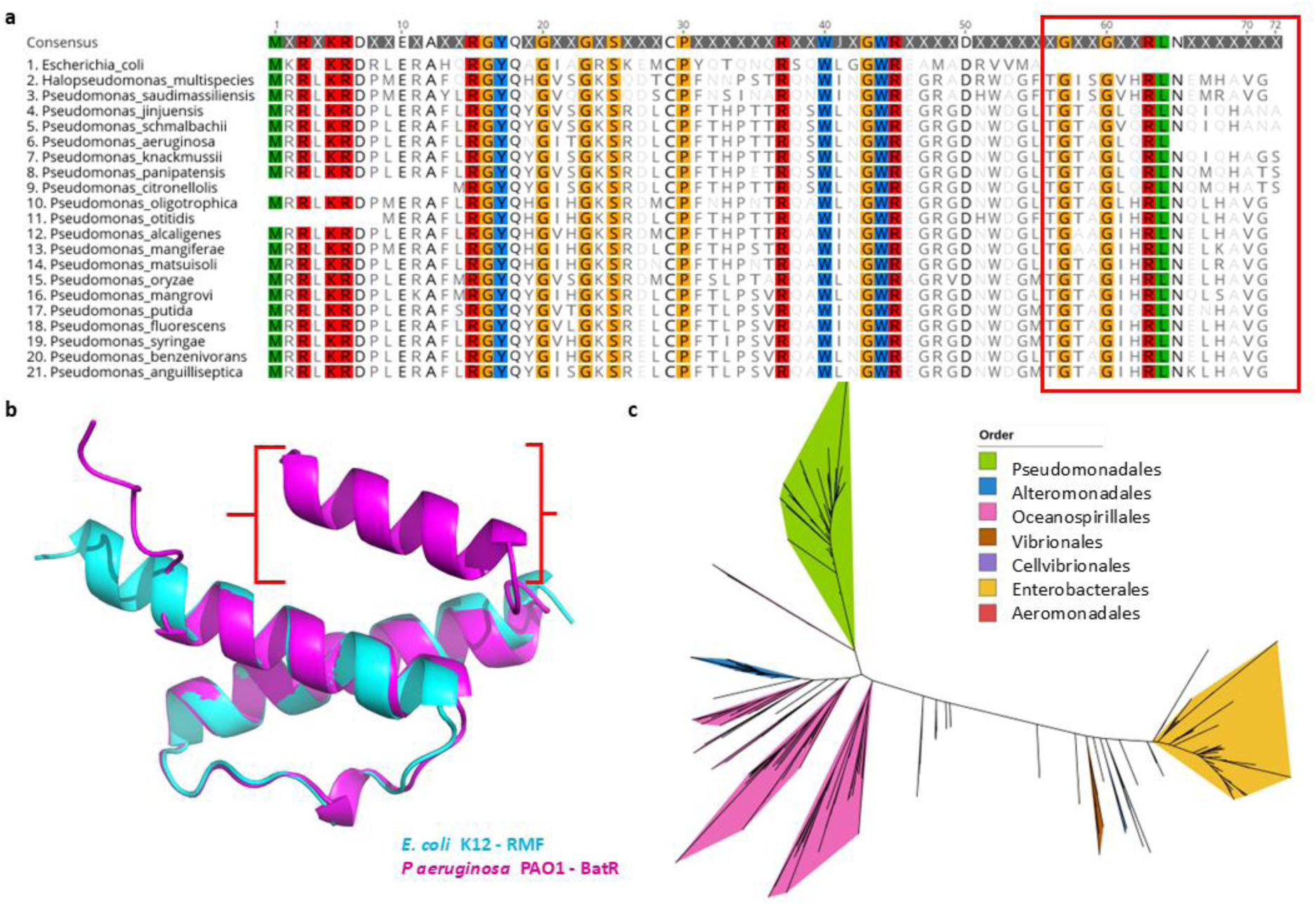
a. **Sequence alignment of BatR and RMF.** ClustalW alignment of RMF from *E. coli* str. K- 12 substr. MG1655 and BatR from 20 different *Pseudomonas* spp. Different colours mark conserved residues in all 21 proteins. The 15-residue C-terminal extension on BatR sequences that is absent from RMF is shown in the red box. **b. AlphaFold 3 model of PAO1 BatR** (magenta), overlaid onto the structure of *E. coli* RMF (cyan). The additional alpha-helix at the C-terminus of the predicted BatR structure is indicated in red. **c. The phylogenetic relationship between BatR/RMF homologs.** The tree is based on 765 proteins homologous to BatR/RMF from γ-proteobacteria.

A phylogenetic analysis of *batR* homologs revealed that these genes are confined to the γ- proteobacteria class (Fig 1c), with distinct evolutionary paths across diverse bacterial orders. Notably, *batR* in Pseudomonadales forms a distinct cluster that diverges significantly from homologs in Enterobacterales, including *E. coli* and *Yersinia pestis*; and Vibrionales, including *Vibrio cholerae,* where RMF plays a role in ribosome hibernation (36). Unlike other characterised RMF genes, *batR* deletion mutants do not exhibit impaired ribosomal integrity during starvation (32). Thus, while *rmf* and *batR* are likely to share an ancestral root, their divergent structure and phylogeny (Fig 1) are consistent with an alternative functional role for BatR (32).

### *batR* protects *P. aeruginosa* biofilms from sub-inhibitory concentrations (SIC) of antibiotics *in vitro*

Given the heightened abundance of *batR* transcripts in dormant *P. aeruginosa* cells within biofilms (12), we assessed the importance of *batR* in biofilm-mediated antimicrobial tolerance *in vitro*. To do this, we generated a non-polar deletion mutant (Δ*batR*) in *P. aeruginosa* PAO1 (hereafter, PAO1) and examined its response to antimicrobial agents when grown under biofilm conditions. Assays were conducted using antibiotics targeting distinct metabolic processes, i.e. β-lactams (piperacillin, PIP) and quinolones (ciprofloxacin, CIP).

The minimal inhibitory concentration (MIC) (37) of the tested antibiotics for cells grown in liquid culture was unaffected in Δ*batR*, (64 µg/mL for piperacillin and 0.5 µg/mL for ciprofloxacin). Interestingly however, Δ*batR* showed a small but significant increase in sensitivity to both PIP and CIP (12-17%), when grown on a solid surface (Fig 2a). We further confirmed the previously observed effect of the aminoglycoside tobramycin (TOB) (12), with Δ*batR* again showing a significant increase in sensitivity on agar (Fig 2a). Next, we used a glass bead biofilm model (38), to assess the survival of cells within established biofilms challenged with PIP. This model revealed a substantial reduction in the survival of established Δ*batR* biofilms compared to WT PAO1, for samples exposed to a sub-inhibitory concentration (SIC) of PIP (Fig 2b). Furthermore, the Δ*batR* phenotype could be fully rescued by the heterologous expression of *batR* (Fig 2b).

**Fig 2.**
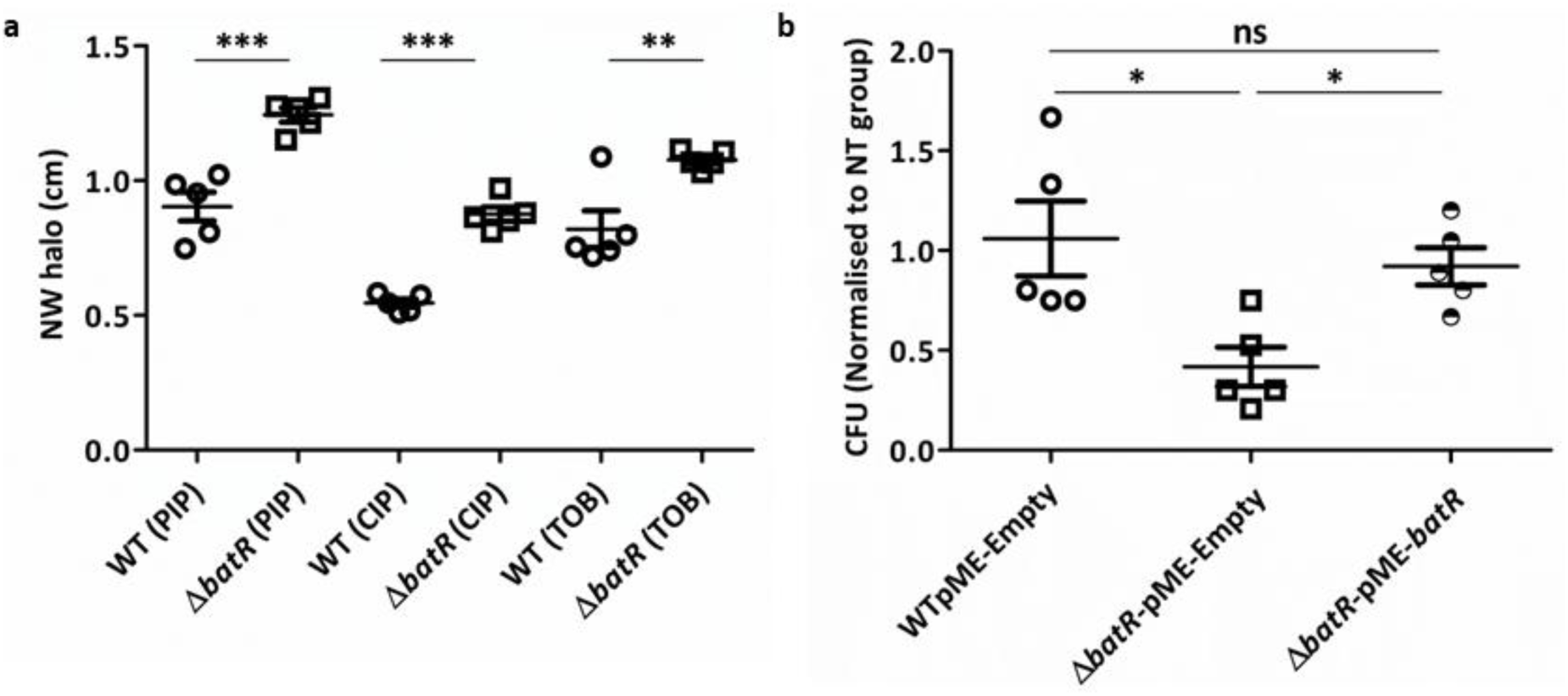
*batR* contributes to biofilm-associated antimicrobial tolerance. a. Antibiotic disk diffusion assay. Results are shown as the normalized width of the antimicrobial halo (NW halo), calculated as described in (39). Results were analysed by a one-way ANOVA and inhibition differed significantly between WT and Δ*batR* strains (F_5,24_= 35.99, p<0.0001). Tukey’s multiple comparisons between WT and Δ*batR* strain under PIP, CIP and TOB treatments are indicated (p<0.001 **, p<0.0001 ***). **b. Glass Beads Biofilm survival**. Bacterial recovery (CFU/bead) from established biofilms grown on glass beads for 24h following treatment with SIC of PIP for 90 min. Strains used were WT PAO1 carrying the empty vector pME6032 (WT-pME-Empty), ΔbatR carrying pME6032 (ΔbatR-pME-Empty), and ΔbatR overexpressing batR (ΔbatR-pME-batR). Normalised values represent the estimated proportion of cells surviving PIP exposure, as described previously (38). Results were analysed by a one-way ANOVA showing significant differences (F2,12=6.414; p<0.05 *). Tukey’s multiple comparisons between WT-pME6032 (PIP) and ΔbatR-pME6032 (PIP), and between ΔbatR-pME6032 (PIP) vs ΔbatR-pME-batR (PIP) are indicated (p<0.05 *).

Since the antibiotics tested in this study have different modes of action, our results suggest that BatR contributes to *P. aeruginosa* antimicrobial tolerance through a general, rather than drug-specific mechanism. A simple explanation was a nonspecific change in membrane permeability. However, we ruled this out by measuring the intracellular concentration of resazurin, a fluorescent dye used to assess permeability and efflux (S1 Fig), which showed little difference between WT and Δ*batR* strains.

### BatR induces specific changes in the PAO1 proteome

To investigate the physiological changes associated with BatR in *P. aeruginosa* biofilms, we conducted a comparative proteomic analysis between the PAO1 WT and Δ*batR* strains. Strains grown on agar plates have been shown to function as organised and coordinated bacterial communities that share multiple characteristics with biofilms (11). Therefore, whole-cell lysates were analysed by TMT proteomics following growth as a lawn on solid medium. On average, 4,581 individual proteins were detected per sample (S1 Data), representing ∼80% of the predicted total *P. aeruginosa* PAO1 proteome (40). Surprisingly, the overall differences between WT and Δ*batR* proteomes were limited (S1 Data), suggesting that the impact of *batR* deletion is quite specific under the conditions tested.

Of the 21 proteins decreased in the Δ*batR* strain (Table 1), seven (PA0617-PA0633) comprise the R2/F2 pyocin gene locus (41). In addition, the loss of *batR* led to reduced abundance of proteins involved in transport & metabolism, two heat shock proteins, and four proteins of unknown function. Conversely, the loss of *batR* increased the abundance of 14 proteins, including Rubredoxin-1, Type VI secretion system components and the Phenazine-1- carboxylate N-methyltransferase PhzM (Table 2).

**Table 1.** COG pathway analysis of proteins decreased in the Δ*batR* strain (TMT proteomics).

| Locus | Gene | Product | COG |
| --- | --- | --- | --- |
| **General function prediction only** |  |  |  |
| PA0617 |  | Probable bacteriophage protein | General function prediction only |
| PA0618 |  | Probable bacteriophage protein | General function prediction only |
| PA0619 |  | Probable bacteriophage protein | General function prediction only |
| <u>PA0620</u> |  | Probable bacteriophage protein | General function prediction only |
| <u>PA0623</u> |  | Probable bacteriophage protein | General function prediction only |
| PA0626 |  | Phage tail protein | General function prediction only |
| PA0633 |  | Phage late control D family protein | General function prediction only |
| PA0803 |  | VOC domain-containing protein | Function unknown |
| <u>PA2853</u> | *oprI* | Major outer membrane lipoprotein (Outer membrane lipoprotein I) | Function unknown |
| PA3796 |  | DUF937 domain-containing protein | Function unknown |
| **Transport & Metabolism** |  |  |  |
| PA3224 |  | 3-phosphoglycerate kinase | Metabolism |
| PA1625 |  | AI-2E family transporter |  |
| PA1906 |  | HIT family protein | Nucleotide transport and metabolism; Carbohydrate transport and metabolism |
| PA2594 |  | PBPb domain-containing protein | Inorganic ion transport and metabolism |
| PA2941 |  | VWFA domain-containing protein | Coenzyme metabolism |
| PA3280 | *oprO* | Porin O | Inorganic ion transport and metabolism |
| PA3113 | *trpF* | N-(5'-phosphoribosyl) anthranilate isomerase (PRAI) | Amino acid transport and metabolism |
| **Translation, Posttranslational modification, protein turnover, chaperones** |  |  |  |
| **PA3049** | ***rmf/batR*** | ***Ribosome modulation factor/ Biofilm antibiotic tolerance regulator*** | ***Translation, ribosomal structure, and biogenesis*** |
| PA5195 | *yrfH* | Heat shock protein 15 | Translation, ribosomal structure, and biogenesis |
| PA4762 | *grpE* | Protein GrpE (HSP-70 cofactor) | Posttranslational modification, protein turnover, chaperones |
| **Cell envelope biogenesis, outer membrane -- Cell motility and secretion** |  |  |  |
| PA1041 |  | Probable outer membrane protein | Cell envelope biogenesis, outer membrane; Cell motility and secretion |

**Table 2.** COG pathway analysis of proteins increased in the Δ*batR* strain (TMT proteomics).

| Locus | Gene | Product | COG |
| --- | --- | --- | --- |

| Energy production and conversion |  |  |  |
| --- | --- | --- | --- |
| PA5351 | *rubA1* | Rubredoxin-1 (RdxS) | Energy production and conversion |
| Lipid Metabolism |  |  |  |
| PA0286 | *desA* | Delta-9 fatty acid desaturase | Lipid metabolism |
| PA3157 | *wbpC* | Probable acetyltransferase | Lipid metabolism |
| Type VI secretion system |  |  |  |
| PA1658 | *hsiC2* | Type VI secretion system contractile sheath large subunit | Intracellular trafficking, secretion, and vesicular transport |
| PA2366 | *hsiC3* | Uricase PuuD | Intracellular trafficking, secretion, and vesicular transport |
| PA2367 | *hcp* | Type VI secretion system tube protein | Intracellular trafficking, secretion, and vesicular transport |
| PA0071 | *tagR1* | FGE-sulfatase domain-containing protein | Hcp secretion island I (HSI-I) type VI secretion system |
| Secondary metabolites biosynthesis, transport, and catabolism |  |  |  |
| PA4209 | *phzM* | Phenazine-1-carboxylate N-methyltransferase | phenazine biosynthetic process |
| PA4143 | *cyaB* | Probable toxin transporter | transmembrane transport |
| PA1436 |  | Probable Resistance-Nodulation-Cell Division (RND) efflux transporter | Defence mechanisms; Inorganic ion transport and metabolism |
| PA2361 |  | IcmF-related_N domain-containing protein | Intracellular trafficking, secretion, and vesicular transport |
| Signal transduction mechanisms |  |  |  |
| PA4154 | *ygiM* | SH3b domain-containing protein | Signal transduction mechanisms |
| Unknown function |  |  |  |
| PA4297 | *tadG* | TadG | Function unknown |
| PA2030 |  | Transmembrane protein | Function unknown |

In *P. aeruginosa* PAO1, the R2/F2 pyocin cluster encodes the structural components for R2 and F2 pyocins, a shared regulatory region and a lytic cassette made up of four proteins (S2a Fig). Among these, the endolysin Lys (PA0629) and the holin (PA0614) are essential for explosive cell lysis, a process in which a subset of cells transition from rod-shaped to round then lyse, releasing intracellular contents including genomic DNA, critical for early biofilm development (24, 42). TMT proteomics revealed increased levels of several R2-type pyocin proteins (PA0617, PA0618, PA0619, PA0620, PA0623 and PA0626) and an F2-type pyocin (PA0633) in the WT strain, suggesting BatR-dependent regulation of the pyocin cluster (S2a Fig). To directly analyse the effect of BatR on explosive lysis, we performed phase-contrast time-lapse microscopy using a microfluidic device (43). We quantified lysis events based on the transition of rod-shaped cells into round morphotypes followed by lysis over a 3h period. Under these conditions, no significant differences in lysis frequencies were observed between WT and Δ*batR* strains (S2b, c Fig). Nonetheless, we cannot rule out a BatR-dependent effect on explosive lysis, as such differences may be undetectable under the low-density, flow-based conditions of the microfluidic setup, but may become more apparent in colony or biofilm contexts, as suggested by the proteomic analysis.

### BatR interacts with a putative stress response kinase

To understand the molecular basis of BatR function, we next investigated its interactions with other PAO1 proteins by performing a co-immunoprecipitation (Co-IP) analysis (S2 Data). BatR- 3xFlag co-immunoprecipitated alongside several cytoplasmic proteins, suggesting potential direct regulatory influences (Table 3). Independent validation of strongly co-precipitating proteins was conducted using Bacterial Two-Hybrid (B2H) analysis. This confirmed specific interaction between BatR and the predicted kinase PA0486, annotated as <u>S</u>tress <u>r</u>esponse <u>k</u>inase A (<u>SrkA</u>) (44)(Figure 3a).

**Fig 3.**
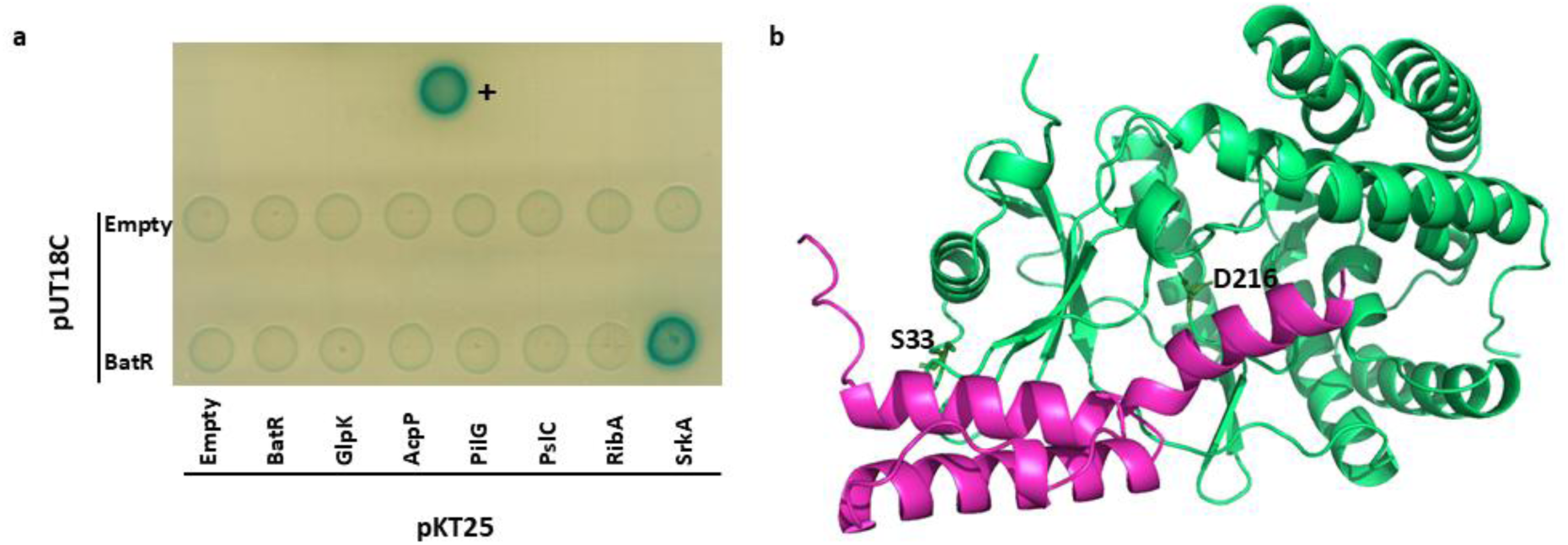
BatR interacts with a predicted kinase protein. **a**. Representative image of qualitative β-galactosidase assays on agar plates. pKT25 and pUT18C fusions are shown in rows and columns, with the indicated protein/empty vector present in each case. Positive control (+): pKT25-zip and pUT18C-zip encoding the two adenylate cyclase fragments, T25 and T18 (45). **b.** Predicted interface between the Alphafold3 models of BatR (magenta) and SrkA (green). The conserved residues Ser33 (S33), predicted ATP-binding site, and Asp216 (D216), predicted Mg^2+^-binding site, in SrkA are shown.

**Table 3.**
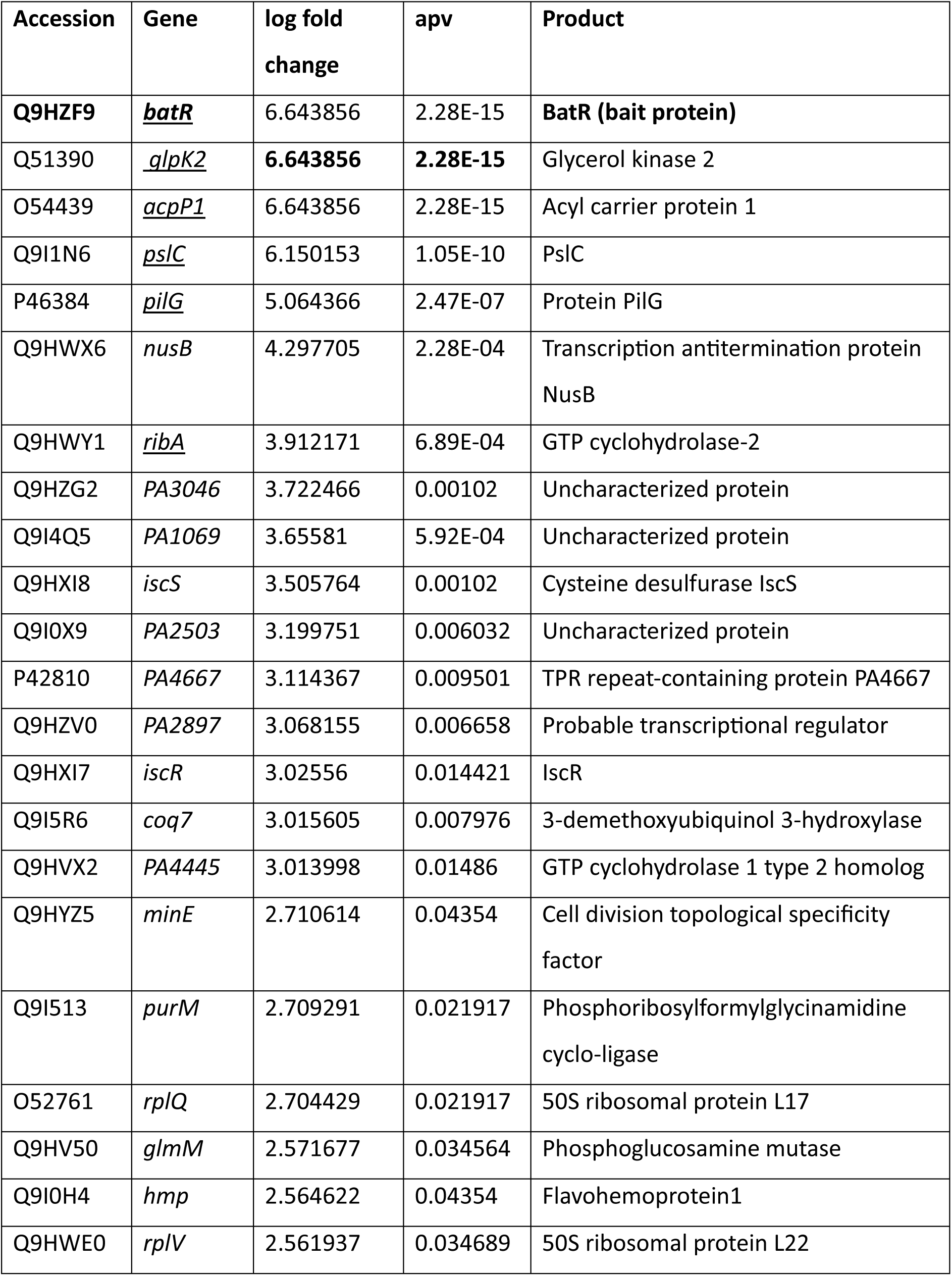
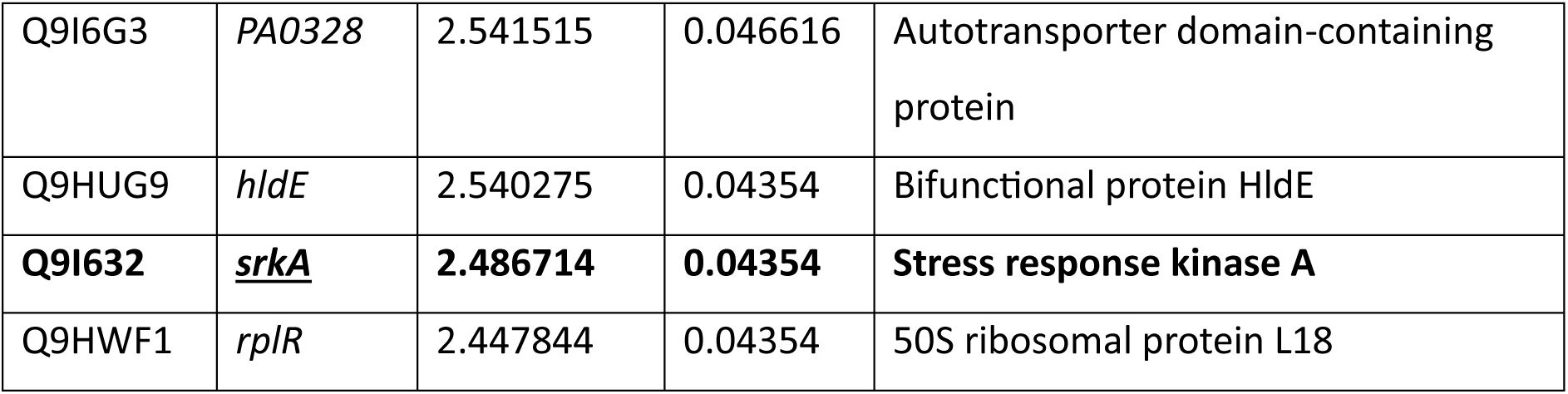
BatR interacting proteins (Co-IP assay).

To gain additional insight into the potential interaction between BatR and SrkA, we used AlphaFold3 (35) to model the potential interactions of BatR with the target protein (Fig 3b). Whilst the Alphafold3 interaction prediction for BatR had low confidence (0.270), the prediction did indicate a potential interaction with the active site of SrkA, including the residue D216, predicted to be the Mg²⁺ binding site.

### SrkA influences biofilm formation; cell death; and pyocyanin production in *P. aeruginosa*

SrkA (PA0486) is annotated as a eukaryotic-like serine-threonine protein kinase. Its homolog in *E. coli* is implicated in protecting cells from stress-induced programmed cell death (44, 46). Since SrkA remains uncharacterized in *P. aeruginosa*, we overexpressed *srkA* in both the WT and Δ*batR* PAO1 strains to assess its functional role. To test whether the observed phenotypes are dependent on SrkA’s kinase activity, we also overexpressed a catalytically inactive (“null”) version of *srkA*, in which the conserved Ser33 and Asp216 residues were substituted with alanine. Alphafold3 analysis of the predicted interaction between BatR and SrkA suggests that BatR binding may interfere with Ser33, which is predicted to be the ATP-binding residue (Fig 3b).

Overexpression of *srkA* led to an increase in biofilm formation in both solid (Congo red binding) and liquid (Crystal Violet staining) media (Fig 4a, b), as well as an increase in pyocyanin production (Fig 4d). This increase in biofilm was independent of the SrkA kinase activity, with the overexpression of the SrkA null mutant showing identical phenotypes to the WT. Overexpression of the kinase-inactive variant also produced a distinctive, highly aggregative colony phenotype on solid media and the emergence of multiple escape mutants, manifesting as regions of reduced dye binding and WT PAO1 morphology (Fig 4a).

**Fig 4.**
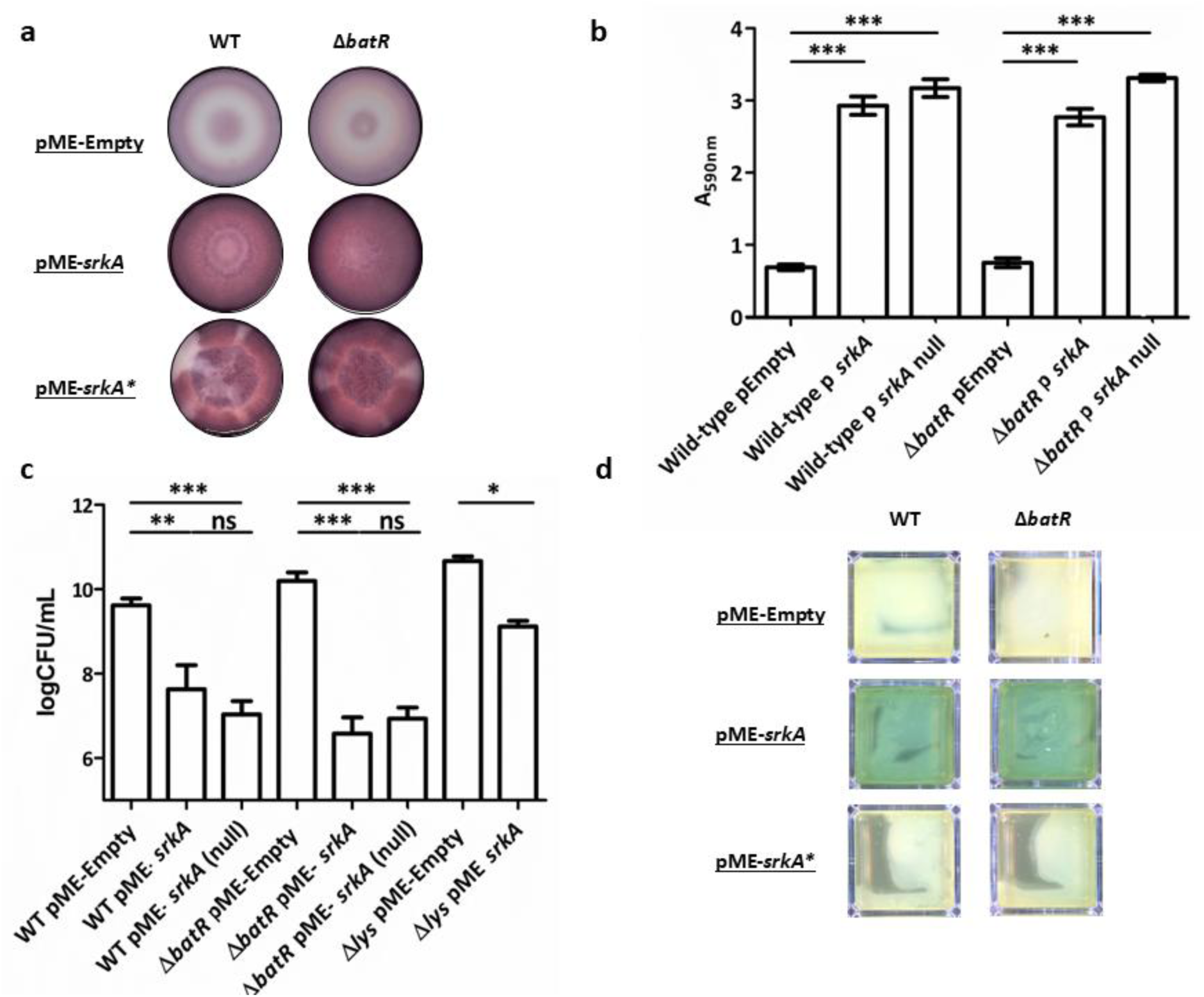
**Phenotypic effects of *srkA* overexpression in PAO1 WT and Δ*batR* strains**. **a. Congo- red binding.** Representative images of WT-pME-Empty, WT-pME-*srkA*, WT-pME-*srkA* null, Δ*batR*-pME-Empty; Δ*batR*-pME-*srkA* and Δ*batR*-pME-*srkA* null biofilms grown using the colony biofilm morphology assay. Only plates supplemented with IPTG are shown (7 days). **b. Biofilm formation.** Strains were grown statically in LB medium for 24h at 37°C. Biofilm biomass was quantified by Crystal Violet staining and measured spectrophotometrically at 590 nm (A_590nm_). Values represent the mean of five biological replicates with two technical replicates each; error bars indicate SD. **c. Cell viability.** Cells were scraped from LB agar plates, resuspended in PBS, and enumerated via serial dilution and plating. **d. Pyocyanin production.** Top view images of cell lawns showing pyocyanin production (blue-green colouration). Results were analysed by a one-way ANOVA showing significance between strains in **b** (F_5,54_= 45.17, p<0.0001) and **c** (F_7,32_= 28.01, p<0.0001). Tukey’s multiple comparisons between strains are indicated (p<0.05 *, p<0.001 **, p<0.0001 ***).

Curiously, the overexpression of *srkA* led to a kinase-independent significant increase in cell death, with even a small induction of SrkA leading to a 10^3^-fold reduction in cell viability. Notably, *batR* appears to confer partial protection against SrkA-mediated killing (Fig 4c).

Given the link between BatR and the R2/F2 pyocin cluster suggested by the proteomic data, we hypothesised that the *srkA*-mediated killing effect occurs through a similar mechanism. To test this, we overexpressed *srkA* in a mutant deficient in production of the endolysin Lys (PA0629) (24). While the impact of *srkA* expression was markedly reduced from that seen in WT PAO1, the Δ*lys* pME-*srkA* mutant still exhibited an approximately 10^2^-fold reduction in viable cell counts compared to the empty vector control (Figure 4c). This suggests that *srkA* induced cell death is partially endolysin-dependent but also implicates an additional lethal pathway that operates independently of the R2/F2 pyocin lysis system.

We next generated a non-polar deletion mutant (Δ*srkA*) and the double mutant (Δ*batR* Δ*srkA*) in *P. aeruginosa* PAO1 and examined their phenotypes. Interestingly, despite SrkA’s dramatic effect when overexpressed, Δ*srkA* showed no phenotypic differences from WT at baseline. However, Δ*batR* Δ*srkA* displayed several strong phenotypes: reduced biofilm formation, increased cell death and increased pyocyanin production (S3 Figure). The fact that these traits emerge only in the absence of both genes suggests that BatR masks or compensates for SrkA- dependent regulatory activities.

### SrkA activates multiple Cell Death mechanisms in *P. aeruginosa*

To visualise the SrkA-mediated killing effect, we repeated the phase-contrast time-lapse microscopy using a microfluidic device. Lysis events increased from 2-3 per 1,000 cells in the control strains (WT and Δ*batR*) (S2 b Fig) to 30-40 per 1,000 cells upon *srkA* overexpression (Fig. 5a). Overexpressing cells appeared highly compromised, with frequent round cells and evidence of explosive lysis (Fig. 5b). Additionally, *srkA* overexpression induced distinct morphological changes, including the emergence of highly elongated filamentous cells (Fig 5b).

**Figure 5.**
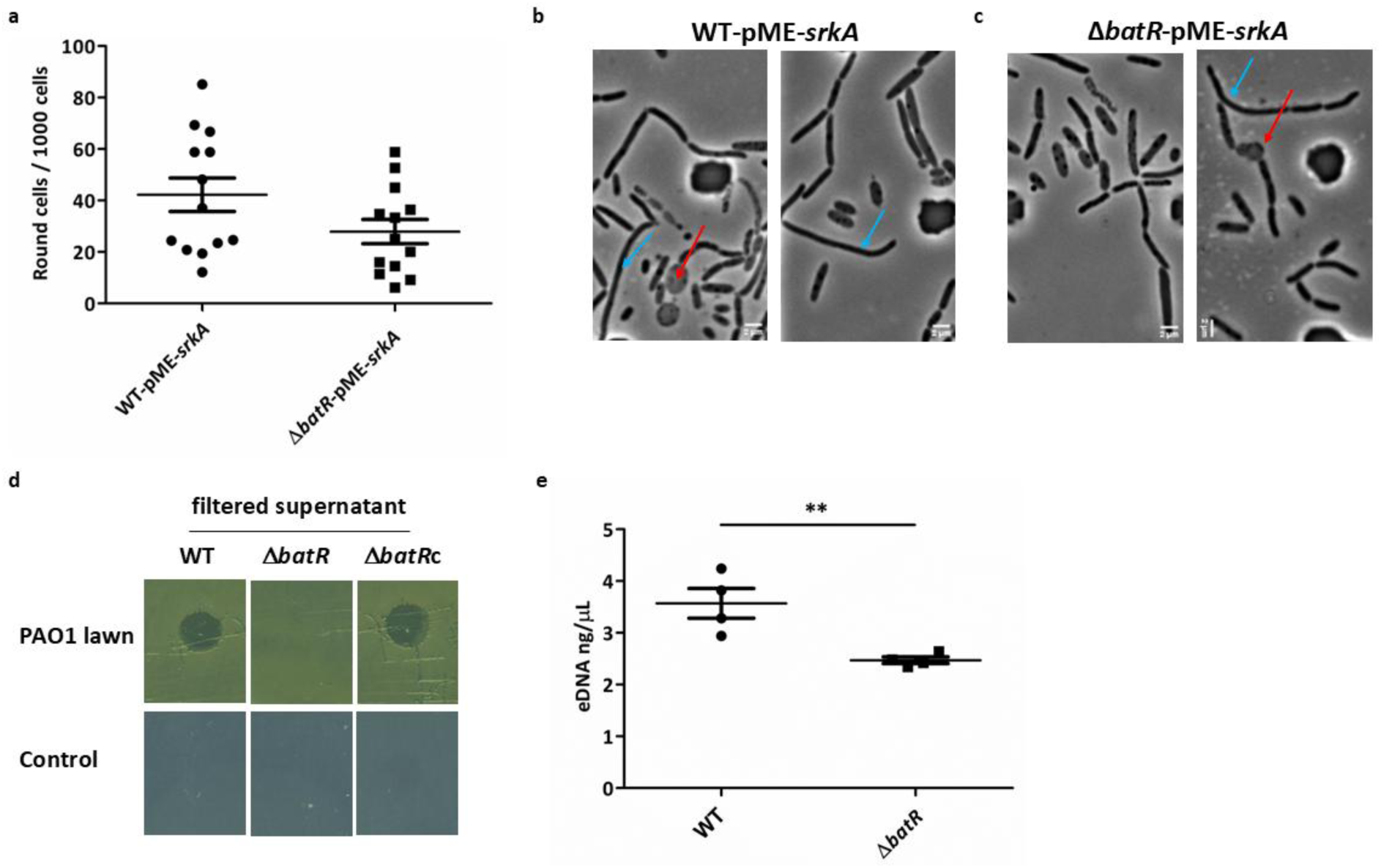
a. ***srkA* overexpression increases explosive cell lysis in *P. aeruginosa*.** Cells were classified as rod-shaped or round over an 11-hour period. The frequency of round cells was calculated as the number of round cells divided by the total number of cells counted at the final time point. No significant differences were observed between strain backgrounds under the conditions tested. **b. *srkA* overexpression alters cell morphology in *P. aeruginosa***. Phase contrast images from an 11h microfluidics assay of WT pME-*srkA* and Δ*batR* pME-*srkA* strains. Cells were incubated for 1h in LB, followed by 9h in LB + 0.1mM IPTG, and a final 1 h in LB. Each experiment was performed independently at least three times. Two primary phenotypes were observed upon *srkA* overexpression: increased frequency of explosive cell lysis (red arrows) and pronounced cell elongation (blue arrows). Time point: 11 h; scale bar: 2 μm. **c. Bacteriophage activity using cell-free supernatants from WT PAO1 and Δ*batR* strains**. **Top:** 20 µL drops from the filtered supernatants of WT, Δ*batR,* and Δ*batR*c (Δ*batR*::*batR*) PAO1 strains were spotted onto the indicator strain, WT PAO1. Zones of clearing on the indicator lawn indicate cell lysis. **Bottom**: a control LB agar plate was spotted with 20 µL of each supernatant to ensure they were free of bacterial growth. Plates were incubated at 37°C overnight. **d. Extracellular DNA quantification**. Double stranded extracellular DNA (eDNA) concentration in the filtered supernatants was quantified on a Qubit Fluorometer using the high sensitivity dsDNA assay kit. The bars denote means of five biological replicates and a T-test shows significant differences between WT and Δ*batR* (t=3.735 df=6, p=0.0097).

To gain further insight into the SrkA mechanism of action, we re-isolated six escape mutants - three from WT PAO1-pME-*srkA* null and three from Δ*batR* PAO1-pME-*srkA* null (Fig 4a) and subjected them to whole-genome sequencing. Comparative analysis of escape mutant genomes revealed that all three mutants derived from the Δ*batR* background had deleted the *srkA* null gene from the overexpression plasmid. In the WT background, one of the three mutants harboured a plasmid-*srkA* null deletion, while a second contained a single-nucleotide polymorphism (SNP) that introduced a premature stop codon at position 147 of the *srkA* null open reading frame, resulting in a truncated and likely nonfunctional protein. The third WT-derived escape mutant sequence harboured multiple unique SNPs in the chromosome, but not in the plasmid, suggesting that bypassing *srkA* null toxicity (without inactivating the gene itself) involves multiple genetic changes. Interestingly, many of these SNPs were in genes encoding oxidoreductases, biofilm-regulators (*gacA, bfmS*)(47), and DNA repair proteins.

We detected a substantial number of SNPs within the PA0717-PA0727 gene cluster, which encodes components of a Pf1-like filamentous bacteriophage (Pf4) in all six strains. This mutational enrichment coincided with increased read depth across the region, consistent with active replication of the prophage (S4 Figure). This observation is notable, as phage Pf4 has previously been implicated in promoting cell death and eDNA release in mature PAO1 and PA14 biofilms (25, 48, 49). Pf4 is unusual among bacteriophages because it lacks a canonical endolysin and is capable of infecting cells within the clonal population from which it originates (48). Furthermore, the highly elongated filamentous cells observed during *srkA* overexpression (Fig 5b), resemble phenotypes associated with Pf4 activity during biofilm development (50). Together, these findings suggest that Pf4 may represent the additional SrkA-triggered lethal pathway that operates independently of the classical R2/F2 pyocin lysis system.

We therefore asked whether BatR also contributes to Pf4 production or activity. To test this, WT and Δ*batR* strains were grown under conditions used for the proteomic assays and filtered culture supernatants were spotted onto lawns of PAO1 WT cells. Interestingly, only WT supernatants produced plaques, whereas Δ*batR* supernatants did not (Figure 5c). This phenotype was fully rescued by heterologous expression of *batR* (Δ*batR*c), implicating BatR in phage-mediated cell death and eDNA release (Figure 5c). Proteomic analysis of these supernatants identified the Pf4 coat protein A among the most enriched proteins in WT samples (Table 4, S3 data). Consistent with this, quantification of eDNA revealed significantly higher levels in WT supernatants compared to those from the Δ*batR* strain (Figure 5d).

**Table 4.** Proteins increased in WT supernatants (Label free proteomics).

| Locus | Product |
| --- | --- |
| PA4831 | Probable transcriptional regulator |
| PA0724 | Coat protein A of bacteriophage Pf1 |
| PA3017 | Universal stress protein |
| PA3901 | Fe(III) dicitrate transport protein FecA |
| PA2795 | tRNA-dihydrouridine synthase A |
| PA5082 | Probable binding protein component of ABC transporter |
| PA0599 | DUF3530 family protein |
| PA1410 | Probable periplasmic spermidine /putrescine-binding protein |
| PA5227 | Cell division protein ZapA (Z ring-associated protein ZapA) |
| PA2800 | VacJ family lipoprotein |
| PA3762 | NGG1p interacting factor NIF3 |
| PA0792 | 2-methylcitrate dehydratase |
| PA4278 | SPOR domain-containing protein |
| PA4447 | Histidinol-phosphate aminotransferase 1 (Imidazole acetol-phosphate transaminase 1) |
| PA0228 | Beta-ketoadipyl-CoA thiolase (3-oxoadipyl-CoA thiolase) |
| PA1339 | Amino acid ABC transporter ATP binding protein |
| PA0862 | TenA family transcriptional regulator |
| PA1674 | GTP cyclohydrolase 1 2 |
| PA3022 | MOSC domain-containing protein |
| PA0729 | Type II toxin-antitoxin system RelE/ParE family toxin |
| PA4741 | 30S ribosomal protein S15 |
| PA2275 | Probable alcohol dehydrogenase (Zn-dependent) |
| PA0052 | Cupin domain-containing protein |
| PA3240 | Putative quercetin 2,3-dioxygenase PA3240 (Putative quercetinase) (Pirin-like protein PA3240) |
| PA0328 | Autotransporter domain-containing protein |
| PA3655 | Elongation factor Ts (EF-Ts) |
| PA4670 | Ribose-phosphate pyrophosphokinase (RPPK) |
| PA4135 | Probable transcriptional regulator |
| PA0904 | Aspartokinase |
| PA1714 | Transcriptional antiactivator ExsD |
| PA0077 | Type VI secretion system component TssM1 |
| PA2082 | Probable transcriptional regulator |
| PA3246 | Pseudouridine synthase RluA |
| PA2756 | 3-isopropylmalate dehydratase large subunit |
| PA0402 | Aspartate carbamoyltransferase |
| PA1204 | NAD(P)H-dependent FMN reductase |
| PA0849 | Thioredoxin reductase |
| PA4653 | Spore coat protein U |
| PA4217 | 5-methylphenazine-1-carboxylate 1-monooxygenase |
| PA4874 | Phosphate starvation-inducible protein PsiF |
| PA0766 | Periplasmic serine endoprotease DegP-like |
| PA4481 | Cell shape-determining protein MreB |
| PA4650 | SCPU domain-containing protein |
| PA1252 | Delta(1)-pyrroline-2-carboxylate reductase |
| PA0660 | NMO domain-containing protein |
| PA0359 | Multidrug transporter |
| PA3997 | Octanoyltransferase (Lipoate-protein ligase B) |
| PA2023 | UTP--glucose-1-phosphate uridylyltransferase |
| PA5520 | Tetratricopeptide repeat protein |
| PA5263 | Argininosuccinate lyase |
| PA0623 | Probable bacteriophage protein |
| PA4847 | Biotin carboxyl carrier protein of acetyl-CoA carboxylase (BCCP) |
| PA1272 | Corrinoid adenosyltransferase |
| PA5257 | HemY_N domain-containing protein |
| PA5558 | ATP synthase subunit b |
| PA0633 | Phage tail protein |
| PA3416 | Probable pyruvate dehydrogenase E1 component, beta chain |
| PA3813 | Iron-sulfur cluster assembly scaffold protein IscU |
| PA1585 | oxoglutarate dehydrogenase (succinyl-transferring) |
| PA3724 | Elastase |
| PA5015 | Pyruvate dehydrogenase E1 component (PDH E1 component) (EC 1.2.4.1) |

### BatR affects the ability of *P. aeruginosa* biofilms to withstand antibiotic challenge in clinically validated infection models

Given BatR’s role in promoting biofilm-mediated antibiotic tolerance under laboratory conditions, we next determined whether this effect extends to clinically relevant infection models. First, we used an *in vitro* synthetic chronic wounds (SCW) model, which mimics the environment of diabetic foot infections (34), to assess biofilm survival following exposure to increasing concentrations of CIP (0-64 µg/mL).

In the absence of antibiotic, no differences in survival were observed between the WT and Δ*batR* strains. However, upon CIP treatment, PAO1 Δ*batR* biofilms showed significantly reduced survival compared to WT biofilms (Fig 6a), with a nearly 40-fold reduction in CFU recovery in the Δ*batR* mutant at higher CIP concentrations.

**Fig 6.**
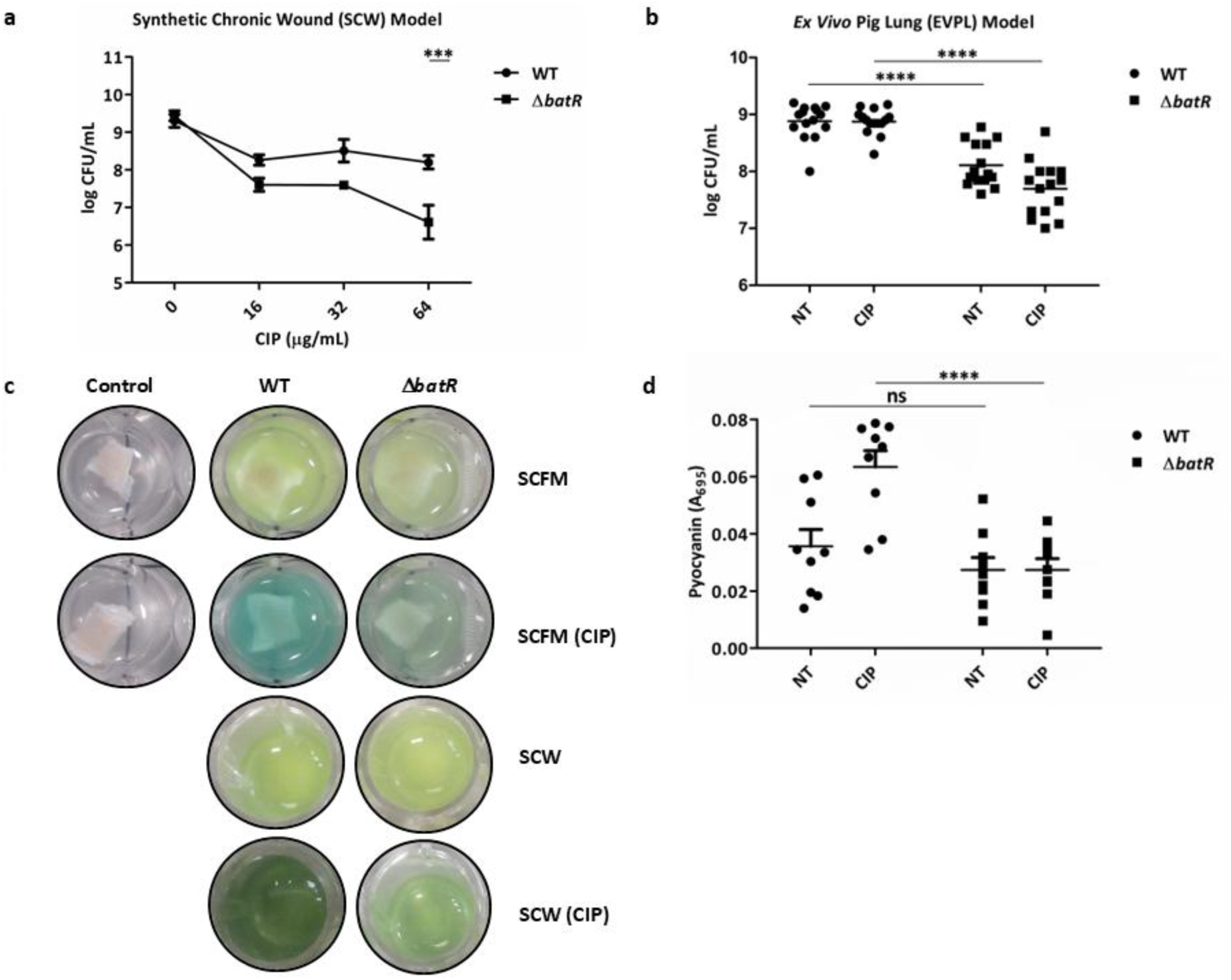
**BatR contributes to antibiotic tolerance and pyocyanin production in clinically validated models. a. Synthetic Chronic Wounds (SCW) Model**. Viability of *P. aeruginosa* PAO1 and the Δ*batR* strain living in established biofilms (24 h), treated with different concentrations of CIP in the cell suspension on top of the matrices. Dots represent the mean from three biological replicates and error bars indicate the standard deviation. Two-way ANOVA showed significant differences for WT and Δ*batR* under 64 µg/mL CIP treatment (p<0.001 **). b. *Ex Vivo* Pig Lung (EVPL) Model. Growth of *P. aeruginosa* PAO1 strain and Δ*batR*, on 15 pieces of EVPL bronchiole (three replicate pieces of tissue from five independent lungs infected per strain) plus Synthetic Cystic Fibrosis Medium (SCFM), with and without 16 µg/mL CIP treatment (17). Colony forming units (CFU) were retrieved from biofilms after 2 d growth at 37°C. Bars denote mean for each genotype across all five lungs, and asterisks denote a significant difference under that condition. Two-way ANOVA analysis showed significant differences for WT vs Δ*batR*; and WT +CIP vs Δ*batR* +CIP (p<0.0001 ***). **c. Biofilm pigmentation.** The characteristic blue-green colouration of *P. aeruginosa* results from a mixture of the exoproducts pyoverdine and pyocyanin. Notably, pigmentation intensity was higher in the WT strain when grown on bronchiolar tissue sections in SCFM and as biofilm aggregates in a collagen gel matrix in the SCW model. b. Quantification of pyocyanin by *P. aeruginosa* WT and Δ*batR* in the EVPL model. Differences in pyocyanin production (A_695_) between WT and Δ*batR* under CIP treatment were significant (p<0.0001***).

To expand these findings to a more complex system, we next used an *ex vivo* pig lung (EVPL) model that recapitulates *P. aeruginosa* biofilm infections in CF bronchioles (33). This model comprises bronchiolar lung tissue and Synthetic Cystic Fibrosis Medium (SCFM) to replicate the luminal mucus environment (51). Biofilms were challenged with CIP at a SIC (16 µg/mL, 1/8^th^ of the MIC in the EVPL model, S5 Fig) and CFUs were enumerated at 2 days postinfection (dpi).

CFU counts in WT biofilms were unaffected by CIP treatment. In contrast, Δ*batR* biofilm survival was reduced even in the absence of CIP, and further decreased upon CIP treatment, resulting in 2.5-fold reduction in bacterial load under antibiotic stress. Our results confirm that BatR contributes to CIP tolerance and supports biofilm establishment within the EVPL model (Fig 6b).

Alongside these effects on biofilm survival, increased blue pigmentation, indicative of pyocyanin production, was evident in WT biofilms compared to Δ*batR* following CIP treatment (Fig 6c, d). A similar effect was also observed in WT biofilms relative to Δ*batR* in the synthetic chronic wound model under antibiotic stress (Fig 6c).

### BatR contributes to biofilm architecture

Next, given BatR’s role in promoting eDNA release under laboratory conditions, we examined the biofilm architecture in infected lung tissue using the EVPL model, with and without SIC of the antibiotic CIP. Infected lung pieces were collected in replica sets and fixed at 2- and 7-dpi, then paraffin-embedded. Sections were stained with H & E (Haematoxylin & Eosin) to visualise total biofilm mass and tissue structure. As expected, distinct biofilm structural differences were observed at both time points (S6 Fig and Fig 7a, b, respectively) between Δ*batR* and WT strains.

**Figure 7.**
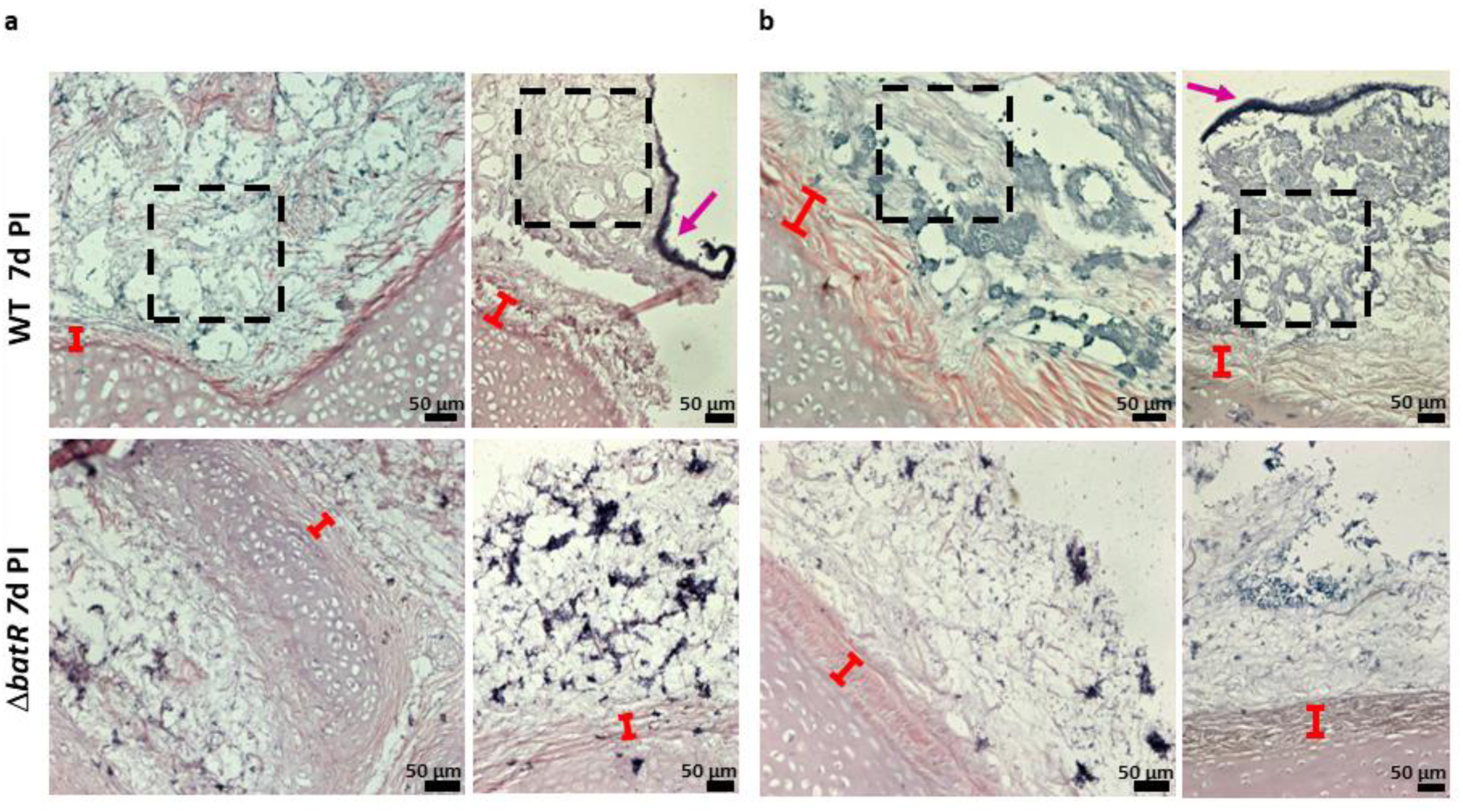
BatR contributes to biofilm architecture. Haematoxylin and eosin (H & E) stained sections of EVPL bronchiolar tissue at 7 dpi. EVPL was infected with *P. aeruginosa* PAO1 WT and Δ*batR*, with uninfected tissue as a negative control. The x20 magnification images from the sections are shown here for **a.** non-treated tissues (SCFM) and **b.** treated with CIP (SCFM + CIP). The cartilage and tissue surface (red bar) stain pink and the bacterial biofilm stain purple, including the bacterial cells and biofilm matrix. The typical sponge-like structure is shown by the dashed boxes on the WT images, and the purple arrow shows the thick layer of matrix covering the biofilm in WT. Representative images of phenotypes at 7 dpi are shown here, but the same results were observed for all biological replicates analysed.

At 2 dpi, the WT strain exhibited significantly higher staining intensity, indicative of denser and more extensive biofilm formation, both in the presence and absence of CIP (S6 Fig). This observation aligns with the higher viability counts observed for the WT strain (Fig 6b), suggesting a larger population of viable cells contributing to the biofilm mass.

By 7 dpi, the EVPL biofilms show a characteristic “sponge-like” appearance consisting of extracellular matrix punctuated by gaps, resembling CF biofilms observed *in vivo* (52–54). Interestingly, qualitative differences in sponge-like architecture were apparent between the Δ*batR* and WT strains, as shown in Fig 7a, b. The WT biofilms showed a thicker matrix and more pronounced structural features, both in untreated samples and those exposed to SIC of CIP. These features were not observed in the Δ*batR* biofilms, suggesting that *batR* influence on biofilm architecture may explain the reduced structural resilience we observe in Δ*batR*, particularly under antibiotic stress (Fig 6b).

## Discussion

In this study we characterise the *P. aeruginosa* proteins BatR and SrkA and determine their contributions to biofilm formation, programmed cell death, pyocyanin production and antimicrobial tolerance, highlighting in turn their potential clinical significance. *batR* homologs exhibit a high degree of conservation within diverse bacteria, particularly in γ-proteobacterial species (36). The identification of a well-supported clade of *Pseudomonadales* carrying *batR* homologs that is both genetically and structurally distinct from the characterised *Enterobacterales*/*Vibrionales rmf* clade suggests a divergent evolutionary path for these genes and supports a distinct cellular function for BatR. The distinct physiological roles of BatR and Rmf showcase how homologous proteins can diverge significantly in function and structure despite sharing elements of sequence conservation, highlighting the importance of experimental validation when inferring protein function across species.

Our findings suggest that *P. aeruginosa* BatR has diverged from its common ancestor with *E. coli* Rmf to regulate phenotypes associated with biofilm formation and stress adaptation during chronic infection. Supporting this, *batR* deletion mutants exhibited a marked difference in biofilm development and an increased sensitivity to multiple antibiotics, including β- lactams and fluoroquinolones, under sessile growth conditions, such as those found in CF and chronic wound infections. Previous transcriptomic analysis of *P. aeruginosa* strain PA14 in EVPL model revealed significant differential expression of *batR* at 7- compared with 1-dpi (51). In addition, *batR* is differentially increased in burn wound infections (55), underscoring the versatility of *batR* in mediating *P. aeruginosa* pathogenesis across various infection settings. eDNA is a critical component of the *P. aeruginosa* biofilm matrix, playing a key role in biofilm development (23). In this study, Δ*batR* formed biofilms in our EVPL model with distinct structural deficiencies compared to those produced by WT PAO1. We speculate that differences observed in biofilm development may be driven in part by differential eDNA release. Our proteomic data support this hypothesis, showing that one-third of the proteins increased in biofilms of the WT strain compared to the Δ*batR* strain are part of the R2-pyocin locus known to induce self-lysis (24).

In addition to its contribution to biofilm development, BatR function appears to be consistently associated with pyocyanin production. Pyocyanin is a critical virulence factor in chronic lung infections in CF patients, impacting gene expression, colony size and biofilm thickness (56). This molecule interacts with O_2_ to form reactive oxygen species, such as hydrogen peroxide (H_2_O_2_), leading to redox imbalance and host cell injury (57). Pyocyanin dependent H_2_O_2_ production also facilitates cell death that contributes to biofilm formation via the release of eDNA (58). In addition, pyocyanin intercalates with eDNA, altering cell surface properties such as hydrophobicity and attractive surface energies, which promotes cell aggregation (59). Within the oxygen-limited environment of *P. aeruginosa* biofilms, pyocyanin is crucial for metabolic continuity and significantly impacts the biofilm’s response to antibiotic treatments (60–62).

Given the hydrophilic nature of BatR (63) and the absence of conserved regulatory domains, we hypothesised that BatR may function by direct interaction with other proteins. This led us to identify an interaction between BatR and PA0486 (SrkA). In *E. coli*, the SrkA kinase has been implicated in stress-induced programmed cell death (44), however its function in *P. aeruginosa* was unknown. We show that in addition to cell death responses mediated via induction of R2/F2 pyocin and bacteriophage Pf4, SrkA positively regulates both biofilm formation and pyocyanin production in *P. aeruginosa*. Curiously SrkA appears to function via two discreet regulatory mechanisms: pyocyanin stimulation requires an active SrkA kinase, whereas the SrkA cell death and biofilm phenotypes were kinase independent.

Programmed cell death is a tightly regulated process typically associated with multicellular organisms, where it promotes organismal fitness by eliminating damaged or unnecessary cells (64, 65). Pyocin and phage-mediated cell lysis fulfil similar roles in bacteria, and can be viewed as a form of bacterial programmed cell death (24). In this context, the biofilm acts as a multicellular community, where the release of public goods such as eDNA benefits the population. Our data suggest that BatR and SrkA modulate three key phenotypes in *P. aeruginosa* PAO1 biofilms: the production of biofilm components, enabling surface attachment and matrix formation; programmed cell death leading to eDNA release; and pyocyanin production, supporting redox chemistry (66). The coordinated deployment of these traits enhances biofilm formation and contributes to population-level resilience.

Our findings support a model in which BatR interacts with SrkA to modulate its activity. As a small protein lacking enzymatic or DNA-binding domains, BatR likely acts as an allosteric partner for other regulators, influencing the activity of SrkA and potentially other proteins under specific conditions. The fact that SrkA-linked phenotypes were only observed in a Δ*srkA* Δ*batR* double mutant supports the existence of a second, as-yet unidentified BatR interaction partner, whose activity compensates for the loss of *srkA* in the single mutant. When both *batR* and *srkA* are missing, this second regulator may trigger the phenotypes we observe in S3 Figure. Identifying additional members of the BatR regulon, alongside the downstream targets of SrkA are high priorities for future research.

*P. aeruginosa* biofilms are a hallmark of chronic infection and mortality in CF patients and are associated with poor clinical outcomes. Mechanisms that regulate susceptibility to cell death within biofilms, such as those mediated by BatR and SrkA, represent potential targets for therapeutic interventions aimed at disrupting biofilm integrity and improving treatment efficacy.

## Materials and methods

### Bioinformatic analysis

A phylogenetic tree of RMF proteins was constructed using 765 publicly available protein sequences from the NCBI database. The dataset was curated based on a criterion of 50% sequence identity and 40% query cover to ensure the representation of diverse homologs while maintaining a reasonable level of similarity. The multiple sequence alignment was performed using Clustal Omega (v1.2.4), generating an alignment matrix with 137 columns and 135 distinct patterns. Phylogenetic signal analysis revealed 82 parsimony-informative sites, 33 singleton sites, and 22 constant sites. Manual curation was conducted to remove repetitive sequences and false hits. The phylogeny estimation was done using IQ-TREE, (multicore v1.6.12), employing the maximum likelihood (ML) criterion. The tree visualization was done using iTOL (v6.8.1), representation chosen was an unrooted tree with branch lengths proportional to the inferred evolutionary distances between sequences. The clade colours were assigned at order level of the taxonomy.

### Bacterial strains and growth media

Bacterial strains and plasmids used in this study are listed in Table 5. Unless otherwise stated, *P. aeruginosa* PAO1 and *E. coli* DH5α strains were routinely cultured in LB (lysogeny broth) (67) at 37°C solidified with 1.5% w/v agar where appropriate. For growth curves, cells were grown at a starting OD_600_ of 0.01 in a 96-well plate. Measurements were taken every 30 min for up to 48 h on a FLUOstar nano plate reader (BMG) with the plate being incubated at 37°C under planktonic conditions.

**Table 5.**
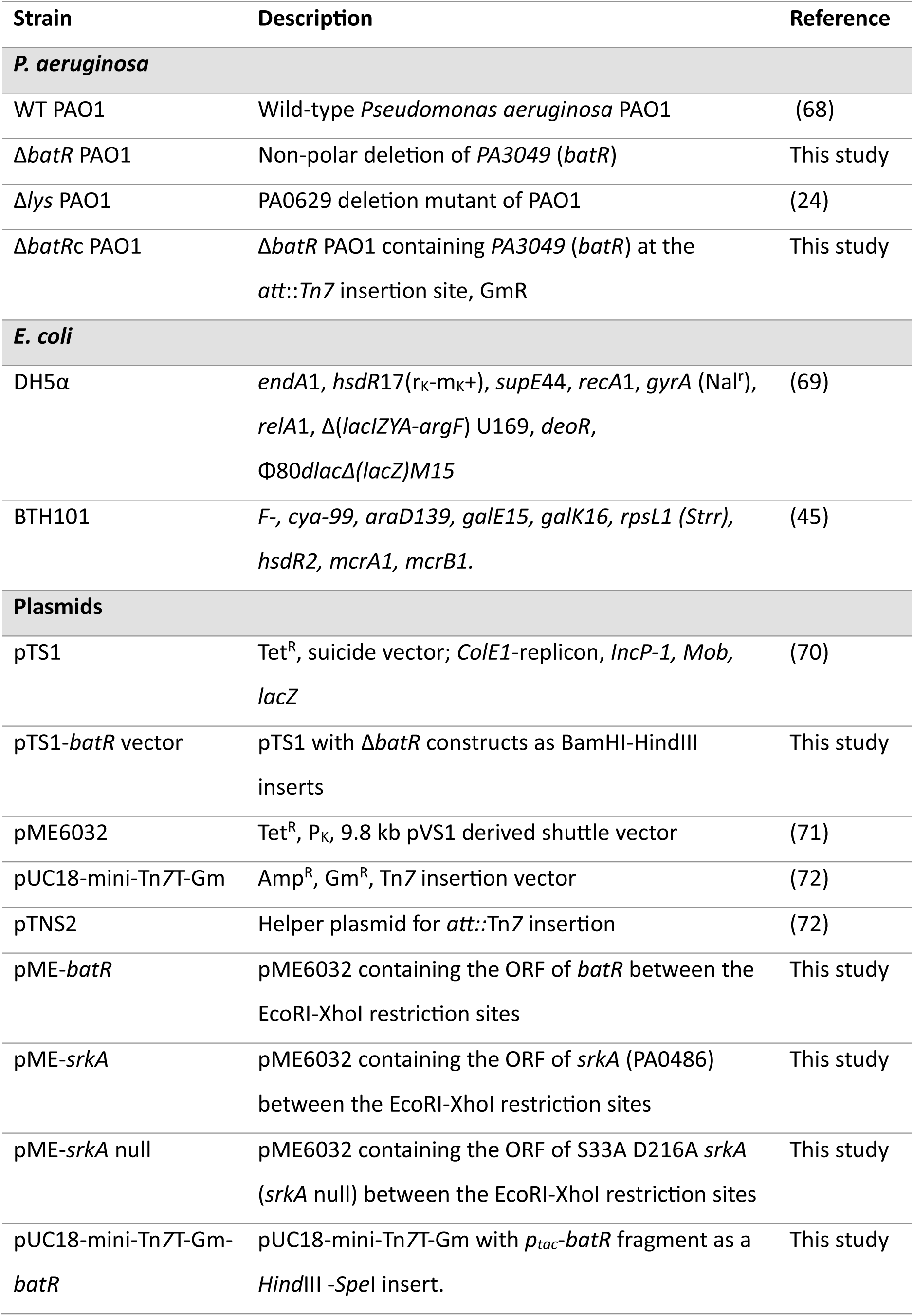
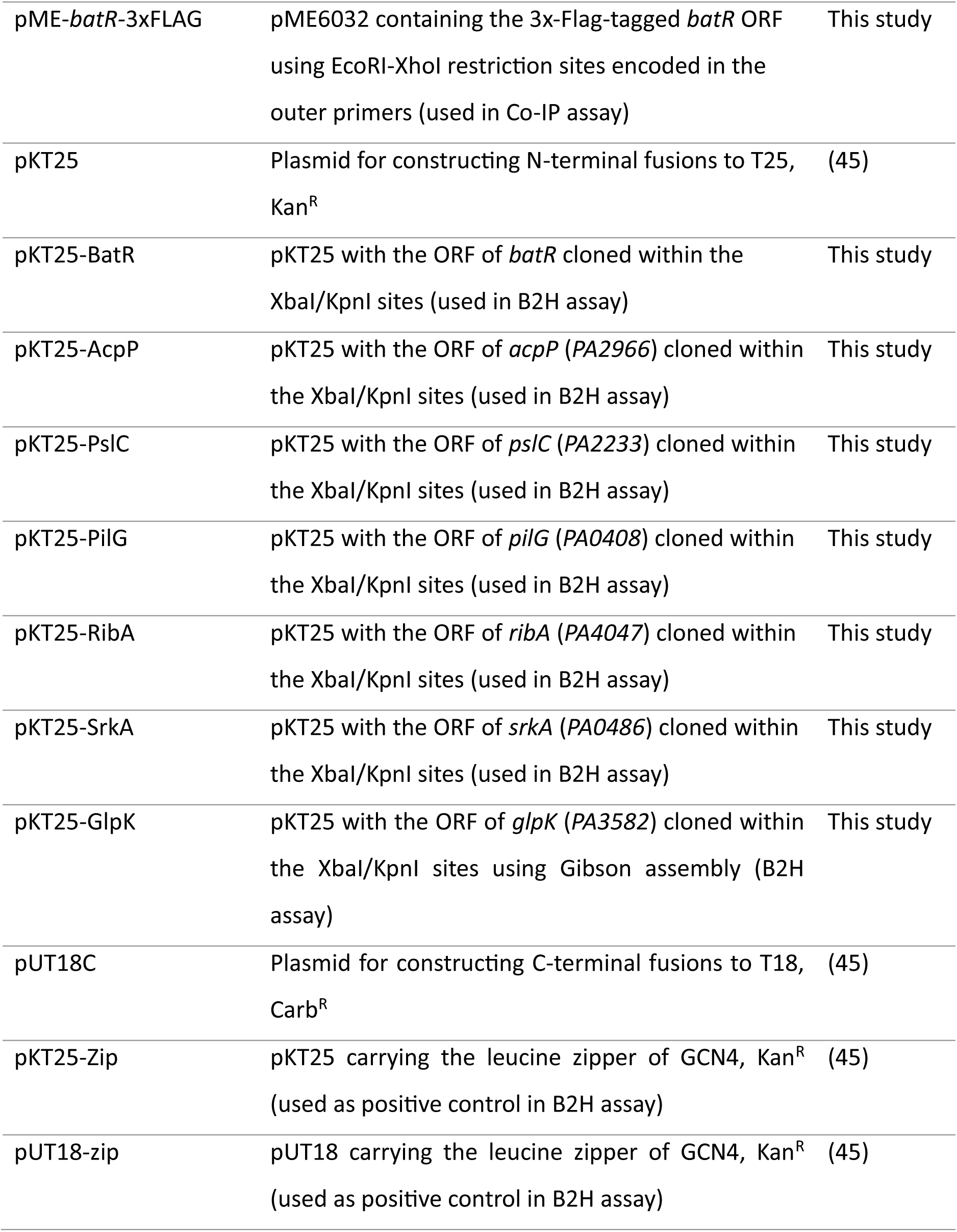
Strains and plasmids used in this study.

Carbenicillin (Carb) was used at 100 μg/mL, Kanamycin (Kan) at 50 μg/mL, Tetracycline (Tet) at 12.5 μg/mL for *E. coli* and 100 μg/mL for *P. aeruginosa*, IPTG at 0.5 mM, and X-gal at 40 μg/mL. The antibiotics Piperacillin (PIP), and Ciprofloxacin (17) were employed at concentrations optimized for each experiment, with details provided accordingly.

### Molecular biology techniques and genetic manipulation of PAO1

These procedures were performed as previously described (73). All pTS1 plasmid inserts were synthesised and cloned into pTS1 by Twist Bioscience.

The ORF of *srkA* with S33 and D216 substituted by an Ala in the *srkA* null mutant was synthesised by Twist Bioscience. The ORFs of *batR* and *srkA* / *srkA* null were amplified by PCR with primers batR_EcoRI_F / batR_XhoI_R and srkA_EcoRI_F / srkA_XhoI_R, respectively (Table 6), and ligated between the EcoRI and XhoI sites of pME6032.

**Table 6.**
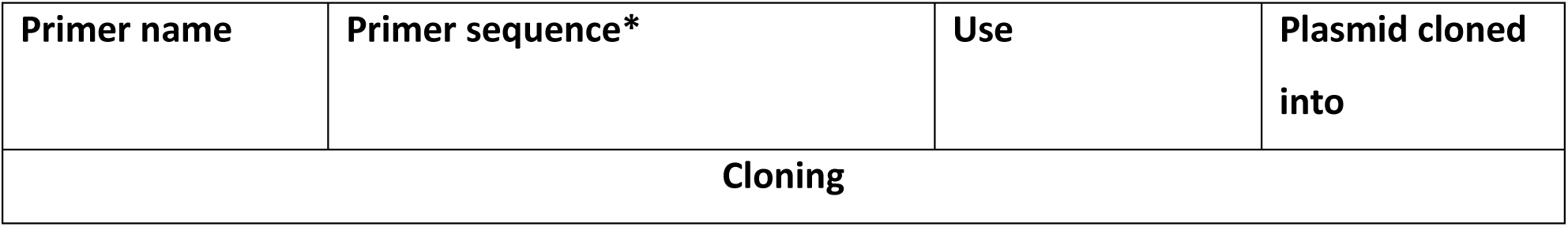
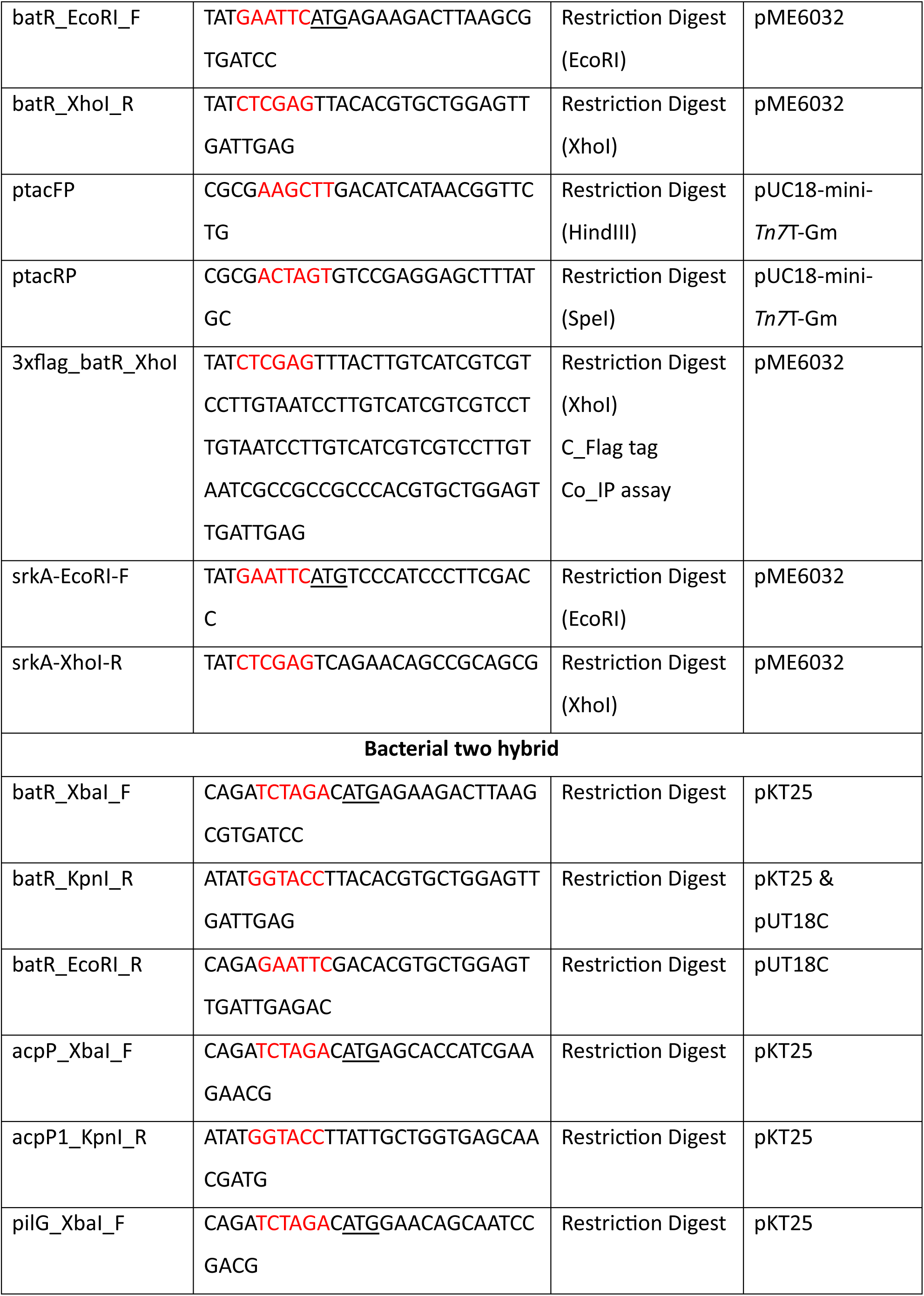
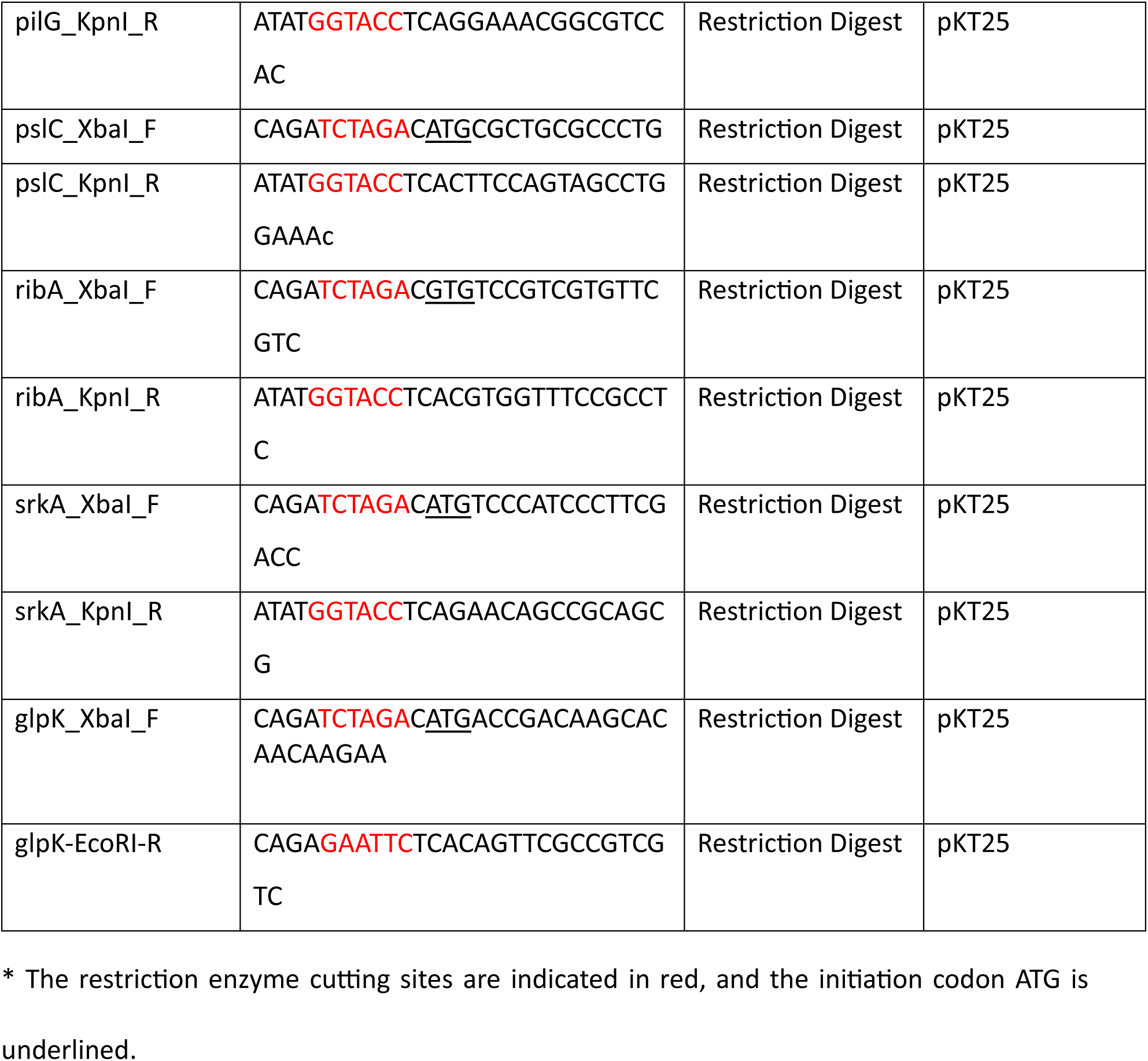
Primers used in this study.

To introduce *batR* gene into the PAO1 *att*::*Tn7* site in the strain Δ*batR*c, the *batR* locus was amplified from the PAO1 genome using primers *batR*_EcoRI_F and *batR*_XhoI_R. The resulting PCR product was cloned into the multiple cloning site of pME6032(71). Primers ptacFP and ptacRP (73) were then used to amplify *batR* in addition to the *tac* promoter and terminator of pME6032. This product was then cloned into the pUC18-mini-*Tn7*T-Gm vector (72) and used to transform PAO1 via electroporation.

For the flag-tagged *batR* construction in pME6032, primers batR_EcoRI_F and 3xflag_batR_XhoI (Table 6) were used. Bacterial-2-hybrid fusion proteins were created by fusing the N-terminus of BatR, AcpP, SrkA, PslC, PilG, GlpK and RibA to pKT25 and combined with pUT18C-Empty and pUT18C-batR (Table 6). Genes were amplified using primers that introduce either XbaI-KpnI (stop codon) or XbaI-EcoRI (stop codon) restriction sites.

### MIC determination

Minimum inhibitory concentrations of antibiotics were determined by the broth microdilution method (37) following the EUCAST guidelines, using Mueller-Hinton broth. The Sub Inhibitory Concentration (SIC) was defined as being 1/8 of the lowest antibiotic concentration that inhibited visible growth after overnight incubation at 37°C.

### Inhibition disc assay

Bacterial cultures were grown in LB medium at 37°C to mid-log phase, A_600nm_=0.5-0.7, and 100 µL were spread on each plate. Whatmann filter paper discs containing the antibiotics were

gently placed on the agar and plates were incubated inverted overnight. The normalized width of the antimicrobial “halo” (NW_halo_) of each disk was determined after (39).

### Glass beads biofilms

These assays were performed as described previously (38, 74) with the following modifications. Ten independent biological replicates were included: five PIP-exposed biofilm lineages (challenged with SIC of PIP for 90 min) and five unexposed control lineages. Cells recovered from the beads were serially diluted and spotted onto LB plates for CFU counting. Results are shown as previously described by (38), where means in PIP-exposed biofilm were then normalised by the average number of cells across all unexposed conditions for plotting. Thus, the values represent the estimated proportion of cells that would survive each exposure for each strain. For complementation assays, culture media were supplemented with 0.5mM IPTG and Tet. The experiment was repeated three times.

### Membrane permeability assays

Differences in membrane permeability to antibiotics were assessed using the resazurin accumulation assay (75). Strains of interest were grown to exponential phase under shaking conditions, using a 1:100 inoculum from overnight cultures. Cells were washed, resuspended in PBS, and normalized for cell density before being mixed with resazurin in round-bottom microtiter plates to a final volume of 100 µL (10 µg/mL resazurin). Fluorescence was measured with an Omega FLUOstar plate reader at an excitation wavelength of 544 nm and an emission wavelength of 590 nm. Five replicates were included per strain, and resazurin-only wells were used as controls. The assay was repeated at least twice, yielding reproducible results each time.

### Quantitative proteomics (TMT) for expression analysis

The experiment was essentially performed as previously described (76) with some modifications, as detailed in Supplementary Information (S4 Data).

### Time-lapse experiment using the CellASIC microfluidic system and image analysis

Time-lapse imaging of *P. aeruginosa* strains was performed as previously described (43), with the following modifications. Briefly, *P. aeruginosa* strains grown overnight in LB medium at 37°C and 250 rpm were sub-cultured to reach an OD_600_ of 0.2. Bacterial cells were loaded into B04A microfluidic plates (ONIX, CellASIC), grown by perfusing LB for 6 h in the case of PAO1 WT and Δ*batR* strains; or perfusing LB for 1 h, then switched to 0.1 mM IPTG LB for 9 h, and LB for 1h in the case of PAO1 strains overexpressing *srkA*. The media flow rate and temperature were maintained at 2 psi and 37°C. Time-lapse imaging was started from the beginning of the experiment and images were acquired every 5 min.

*P. aeruginosa* strains were visualized using a Zeiss Axio Observer Z.1 inverted epifluorescence microscope fitted with a sCMOS camera (Hamamatsu Orca FLASH 4), a Zeiss Colibri 7 LED light source, a Hamamatsu Orca Flash 4.0v3 sCMOS camera, and a temperature-controlled incubation chamber. Images were acquired using a Zeiss Plan Apochromat 100x/NA 1.4 Ph3 objective. Still images and time-lapse images series were collected using Zen Blue (Zeiss) and analysed using Fiji (77).

### Co-immunoprecipitation and mass spectrometry analysis

This protocol is detailed in Supplementary information (S5 Data).

### Bacterial 2 hybrid assays

These assays were performed as described elsewhere (76) with some modifications. The ORFs of *batR, acpP, pilG, pslC, ribA, srkA* and *glpK* were cloned into pKT25, and pUT18C using either conventional restriction enzyme cloning or Gibson assembly, as indicated in Table 6.

### Alphafold3 model predictions

Protein structures were predicted from amino acid sequences using the Alphafold3 server (alphafoldserver.com). For PAO1 BatR (AF_AFQ9HZF9F1) and *E. coli* Rmf (AF-P0AFW2-F1-v4), initially protein structures were retrieved from AlphaFoldDB with interactions modelled independently using the AlphaFold3 server. For interactions, both the bait (BatR WT) and target (SrkA) amino acid sequences were input into the sever and predictions were modelled using default settings. Models were ranked based on their respective PTM/iPTM score and the highest ranked score was taken forward for analysis. Structural analysis and alignments were conducted in Pymol.

### Colony biofilm morphology assay

Agar plates for colony morphology experiments were prepared as previously described (78). Briefly, a mixture of 1% agar and 1% tryptone was autoclaved and cooled to 60°C before 20 μg/ml Coomassie blue, 40 μg/ml Congo red, Tetracycline, and 0.05mM IPTG were added. For colony spotting, five microliters of cultures at an OD_600_ of 0.5 were spotted on plates and incubated for up to 7 days at 23 to 25°C.

### Cell viability and pyocyanin production phenotypes

Five biological replicates of each strain were grown in LB medium at 37°C to mid-log phase (A_600nm_=0.2-0.3). Then, 25 µL of each culture were spread onto wells of 24-well plates. When required, tetracycline and IPTG were added at the appropriate concentrations. Plates were incubated for 24h at 37°C and scanned. For cell viability measurements, cells were scraped from the agar and resuspended in 1mL of 1X PBS. These resuspensions were serially diluted and spotted onto LB agar plates for CFU enumeration.

### Crystal Violet (CV) assays

For the CV assay, five biological replicates of each strain were grown overnight in LB medium supplemented with Tet and then diluted into 200 μL of fresh LB to give an OD_600nm_ of 0.50 in microtiter plates. The expression of recombinant PAO1 *srkA*, was induced with 0.05mM IPTG. The microtiter plates were incubated at 37°C for 24 h, after which the wells were emptied and rinsed three times with water before staining. For staining, 200 μL of 0.1% CV was added to each well and incubated for 15 min at room temperature. The crystal violet dye was then removed, and the wells were rinsed with water. The dye bound to the cells was then dissolved in 70% ethanol, and the A_590nm_ was measured using a SPECTROstar nano plate reader (BMG Labtech).

### Genomic analysis

Six escape mutants -three from WT PAO1-pME-*srkA* null and three from Δ*batR* PAO1-pME- *srkA* null- were chosen for whole-genome sequencing. These colonies were re isolated from the colony biofilm morphology assay at day 7. As controls, the genomes of WT and Δ*batR* PAO1 strains were also sequenced. Whole-genome sequencing was performed by Plasmidsaurus using the Oxford Nanopore long-read technology. Variants were called against the ancestral reference genome using the Breseq computational pipeline using the polymorphic settings (79). All variants were validated visually using the alignment viewer IGB (80).

### Bacteriophage experiments

Strains of interest were grown overnight in 10 mL of LB broth at 37°C. Cultures were adjusted to an OD_600_ of 1.0, and 100 µL of the normalized cultures were spread onto LB agar plates. Plates were incubated for 24 hours at 37°C and the resulting bacterial lawns were scraped from the plates and resuspended in 5 mL of 1 X PBS. These homogenates were passed through a 0.2 μm syringe filter to remove bacterial cells.

Indicator strains were grown overnight in 10 mL of LB broth and diluted 1,000-fold in PBS. Then, 50 µL of the diluted indicator cultures was spread evenly onto LB agar plates. Next, 10 µL of the filtered supernatants was spotted onto the indicator strain plates. Plates were incubated overnight at 37°C. As a control, filtered supernatants were spotted onto LB agar plates to confirm that they were cell-free.

### Label free proteomics

This protocol is detailed in Supplementary information (S6 Data).

### Synthetic chronic wound infection model

These assays were performed as previously described (34). Briefly, batch cultures were grown at 37°C in Synthetic Wound Fluid (SWF) overnight to early mid log phase (approx. 6 h). The synthetic wounds were prepared as follows: For 10 mL collagen solution (2 mg/mL), 1 mL 0.1% acetic acid was mixed with 2 mL collagen stock solution (10 mg/mL) and kept on ice. Then, 6.0 mL of cold Synthetic Wound Fluid (SWF) (50% foetal bovine serum (Gibco 10270) and 50% Peptone water (Fluka 70179)) was added followed by 1 mL 0.1 M NaOH. After mixing, 200 μL of the collagen solution was added to each well (24-well plates). To achieve a complete polymerization of the collagen, the plates were placed in an incubator at 37°C for 1 h.

The starter bacterial cultures were diluted at an OD_600_ of approx. 0.05 – 0.1 in SWF. For small wounds, 50 μL of the diluted starter culture was added to each synthetic wound. Plates were incubated at 37°C for 24 h. After this time, 100 μL of antibiotic CIP at different concentrations was added to the wounds and plates were returned to 37°C for a further 24h. For bacterial recovery, 0.5 mg/mL collagenase (made in PBS) was added to each wound and incubated at 37°C for 1 h. Serial dilutions of these homogenates were plated on LB agar for CFU counting.

### EVPL infection model

EVPL was prepared as previously described (51, 81). Briefly, porcine lungs were obtained from two local butchers (Quigley and Sons, Cubbington and Taylor’s Butcher, Earlsdon) and dissected on the day of delivery under sterile conditions. The pleura of the ventral surface was heat sterilised using a hot pallet knife. A sterile razor blade was then used to make an incision in the lung, exposing the bronchiole. A section of the bronchiole was extracted, and the exterior alveolar tissue removed using dissection scissors. Bronchiolar sections were washed once in a 1:1 mix of Dulbecco’s modified Eagle medium (DMEM) and RPMI 1640 supplemented with 50 μg/mL ampicillin (Sigma-Aldrich) then cut into approximately 5 mm wide longitudinal strips. The bronchiolar strips were placed in a second 1:1 DMEM, RPMI 1640 supplemented with 50 μg/mL ampicillin wash and cut into squares approximately 5 x 5 mm in size. The tissue squares were washed for a third time in 1:1 DMEM, RPMI 1640 containing 50 μg/mL ampicillin. Bronchiolar pieces were then further washed in SCFM, UV sterilised for 5 min and transferred to individual wells of a 24-well plate containing 400 μL SCFM supplemented with 20 μg/mL ampicillin and solidified with 0.8% (w/v) agarose per well.

A sterile 29G hypodermic needle (Becton Dickinson Medical) was touched to the surface of a *P. aeruginosa* colony grown on LB agar overnight at 37°C and used to pierce an individual piece of bronchiolar tissue. Uninfected control tissue sections were mock inoculated with a fresh, sterile needle. Following infection of bronchiolar tissue pieces, 500 μL of SCFM ± 16 μg/mL CIP was added to each well. Tissue pieces were incubated at 37°C with a Breathe-Easier® membrane (Diversified Biotech) for 2 and 7 d.

### EVPL biofilm recovery, assessment of bacterial load and pyocyanin production

EVPL biofilm recovery and assessment of bacterial load and virulence factor production were determined as described (51). Briefly, bronchiolar tissue pieces were removed from the 24- well plate following incubation, and each briefly washed in 500 μL PBS in a fresh 24-well plate to remove planktonic cells. Tissue pieces were then transferred into sterile homogenisation tubes (Fisherbrand) containing eighteen 2.38 mm metal beads (Fisherbrand) and 1 mL PBS. Tissue was bead beaten in a FastPrep-24 5G (MP Biomedicals) for 40 s at 4 m/s to recover the bacteria and virulence factors from the tissue-associated biofilm. To determine the bacterial load, the homogenate was serially diluted in PBS and plated on LB agar. Plates were incubated overnight at 37°C and colony counts used to calculate colony forming units (CFU) per tissue piece.

To quantify pyocyanin produced by the *P. aeruginosa* biofilms, the homogenate was diluted 1:4 in PBS to obtain a sufficient volume for further experiments. The diluted homogenate was passed through a 0.2 μm filter to remove bacterial cells and tissue debris. Total pyocyanin was quantified by measuring absorbance of homogenates at 695 nm (33).

### Haematoxylin & eosin staining

*P. aeruginosa* infected EVPL tissue pieces and uninfected control tissue were fixed in 10% (v/v) neutral buffered formalin (VWR Chemicals), as previously described (51). The infected/uninfected EVPL tissue pieces were fixed and sent to the University of Manchester’s Histology Core Facility for paraffin wax embedding, sectioning, and mounting. Mounted tissue sections were de-paraffinized in xylene for 20 min. To re-hydrate the tissue, slides were transferred to 95% (v/v) ethanol followed by 70% (v/v) ethanol. Any residual ethanol was removed by washing slides in distilled water. Samples were stained in Mayer’s hemalum solution (Merck Millipore) then washed in running tap water (1 L tap water with approximately 10 g sodium carbonate). Samples were counterstained in eosin Y solution (Merck Millipore) for then dehydrated by dipping the slides in 95% (v/v) ethanol. Samples were transferred to fresh 95% (v/v) ethanol then placed in 100% (v/v) ethanol. The samples were then placed in xylene. The samples were mounted using DPX mounting fluid and images were taken using a Zeiss Axio Imager Z2 light microscope with the Zeiss AxioCam 506 and Zeiss Zen Blue v2.3 pro software.

### Data presentation and statistical analyses

All graphs and statistical analyses, including one-way and two-way ANOVA followed by post-hoc Tukey’s Multiple Comparison Test, when appropriate, were performed using GraphPad Prism version 5.04 for Windows.

## Data availability

The TMT proteomics data have been deposited to the ProteomeXchange Consortium via the PRIDE (82) partner repository with the dataset identifier PXD050997 and 10.6019/PXD050997.

The label-free proteomic data have been deposited to the ProteomeXchange Consortium via the PRIDE (85) partner repository with the dataset identifier PXD062857 and 10.6019/PXD062857.

Co-IP data are available via ProteomeXchange with identifier PXD050995.

## Acknowledgments

The authors would like to thank Masanori Toyofuku and Nobuhiko Nomura for the strain Δ*lys* PAO1, and Jovana Kaljevic for her support with the microfluidics experiments and image analysis.

## Author contributions

A.P. and J.G.M. conceptualized the project. A.P. performed all the experiments and data analysis presented here, except for the following assays: C.M.A.T. performed AlphaFold modelling and associated analyses; G.C. performed the genome sequencing and the proteomic analyses; H.D. performed the phylogenetic analysis and sequence alignment of *batR* homologs; E.T. carried out the membrane permeability assay; G.S. and C.M. performed TMT and label-free proteomics, the Co-IP assay, and their respective data analyses.

A.P., F.H., and J.G.M. contributed to funding acquisition, supervision and project management. M.A.W. and F.H. provided valuable feedback on the clinical significance of the experiments. A.P. wrote the full manuscript, and all authors contributed to the final version.

---

**S1 Fig.**
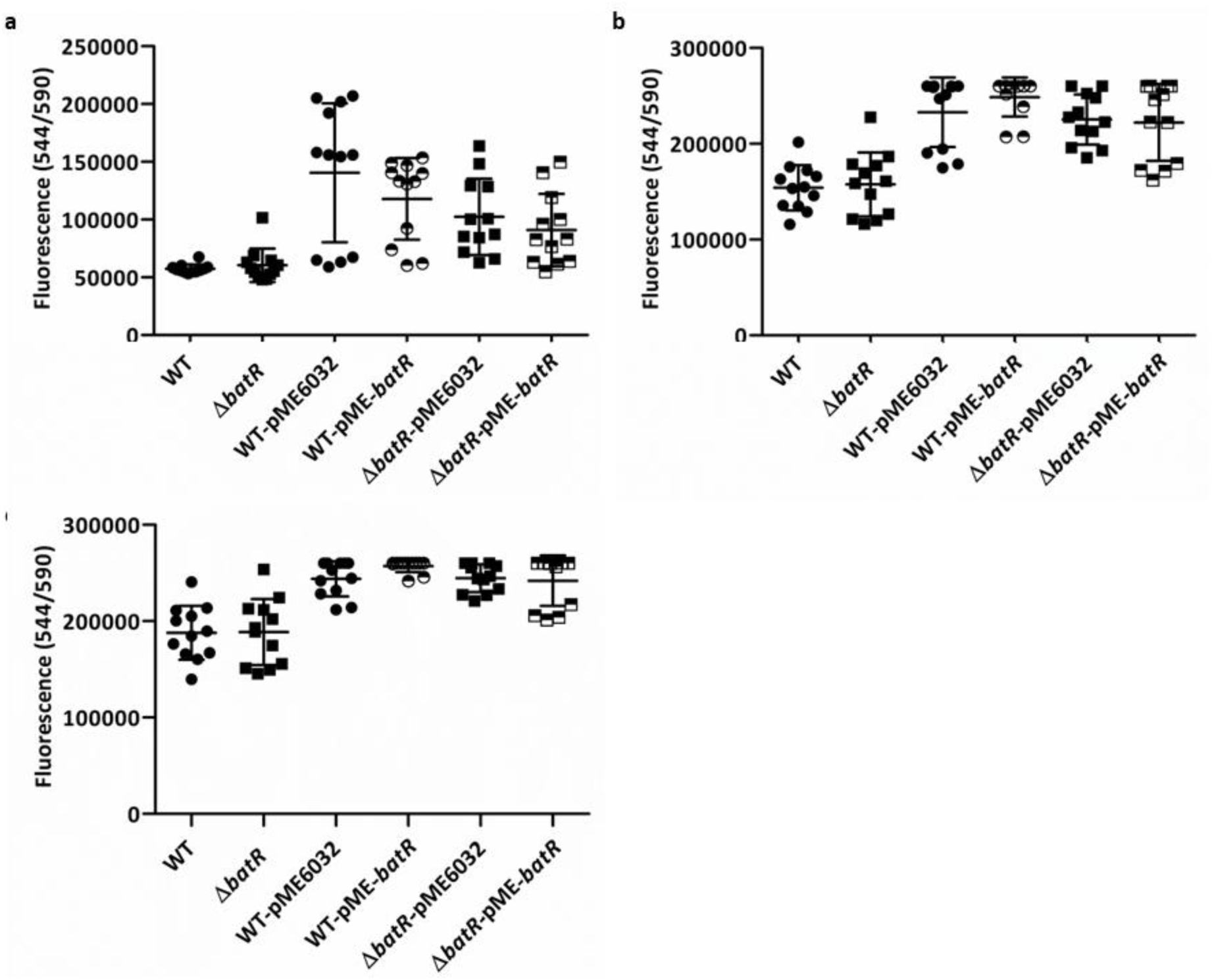
BatR does not affect membrane permeability. Drug accumulation was assessed by measuring resazurin fluorescence at an excitation wavelength of 544 nm and an emission wavelength of 590 nm (544/590 nm) for 2 hours (a), 12 hours (b) and 18 hours (c). Lines represent the mean of three biological replicates, each with four technical replicates, with the error bars showing the standard deviation. No significant differences were observed between the strains tested.

**S2 Fig.**
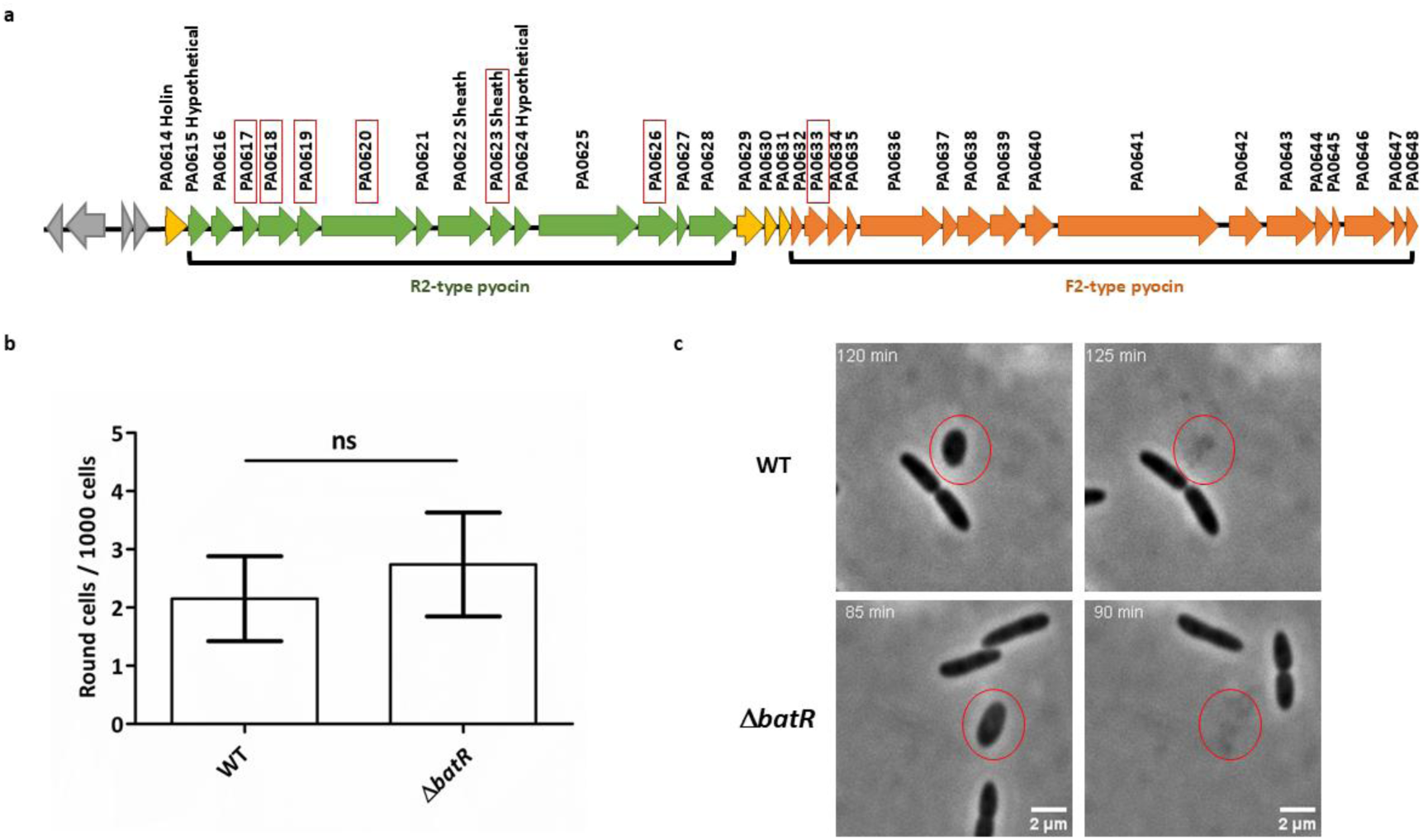
a. Genetic organization of the R2 pyocin gene locus in *P. aeruginosa* PAO1. Structure of the R2-F2 pyocin gene cluster in *P. aeruginosa* PAO1. Grey indicates regulatory genes; yellow, the lysis cassette; green, R2-type pyocin genes; and orange, F2-type pyocin genes. The function of individual proteins is labelled, and proteins enriched in the WT strain proteome are boxed. Gene sizes are drawn to scale. **b. Proportion of cells exhibiting a round-cell morphotype in microfluidics assays.** Cells were classified as either rod-shaped or round over a 3-hour period, and the frequency of round cells was calculated as the number of round cells divided by the total number of cells counted at *t* = 180 min. Results were analysed by a one-way ANOVA showing no significant differences between WT and Δ*batR* strains under the conditions tested. **c.** Phase-contrast time-lapse images showing rod-to-round cell transition and explosive lysis events (highlighted by red circles) in WT and Δ*batR* strains. Time is indicated in minutes (top right); scale bar, 2 μm.

**S3 Figure.**
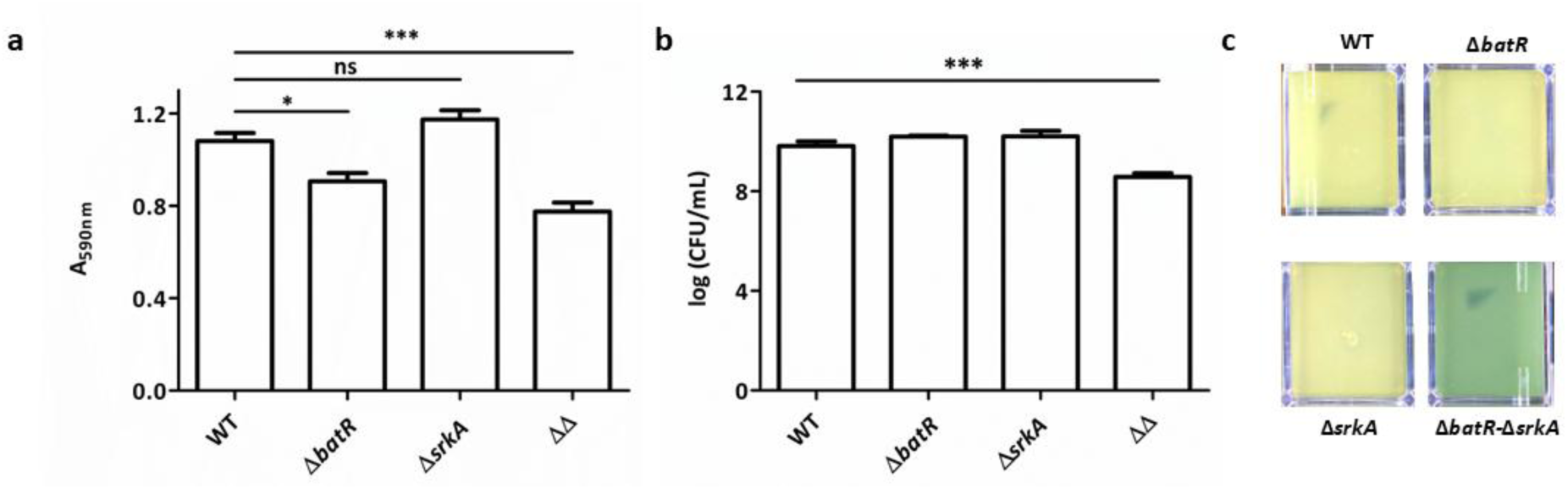
Phenotypes of PAO1 WT; Δ*batR;* Δ*srkA;* and Δ*batR* Δ*srkA* strains. a. Biofilm formation. Strains were grown statically in LB medium for 24h at 37°C. Biofilm biomass was quantified by Crystal Violet staining and measured spectrophotometrically at 590 nm (A_590nm_.). Values represent the mean of five biological replicates with two technical replicates each; error bars indicate SD. **b. Cell viability.** Cells were scraped from LB agar plates, resuspended in PBS, and enumerated via serial dilution and plating. **c. Pyocyanin production.** Top view images of cell lawns showing pyocyanin production (blue colouration).

**S4 Figure.**
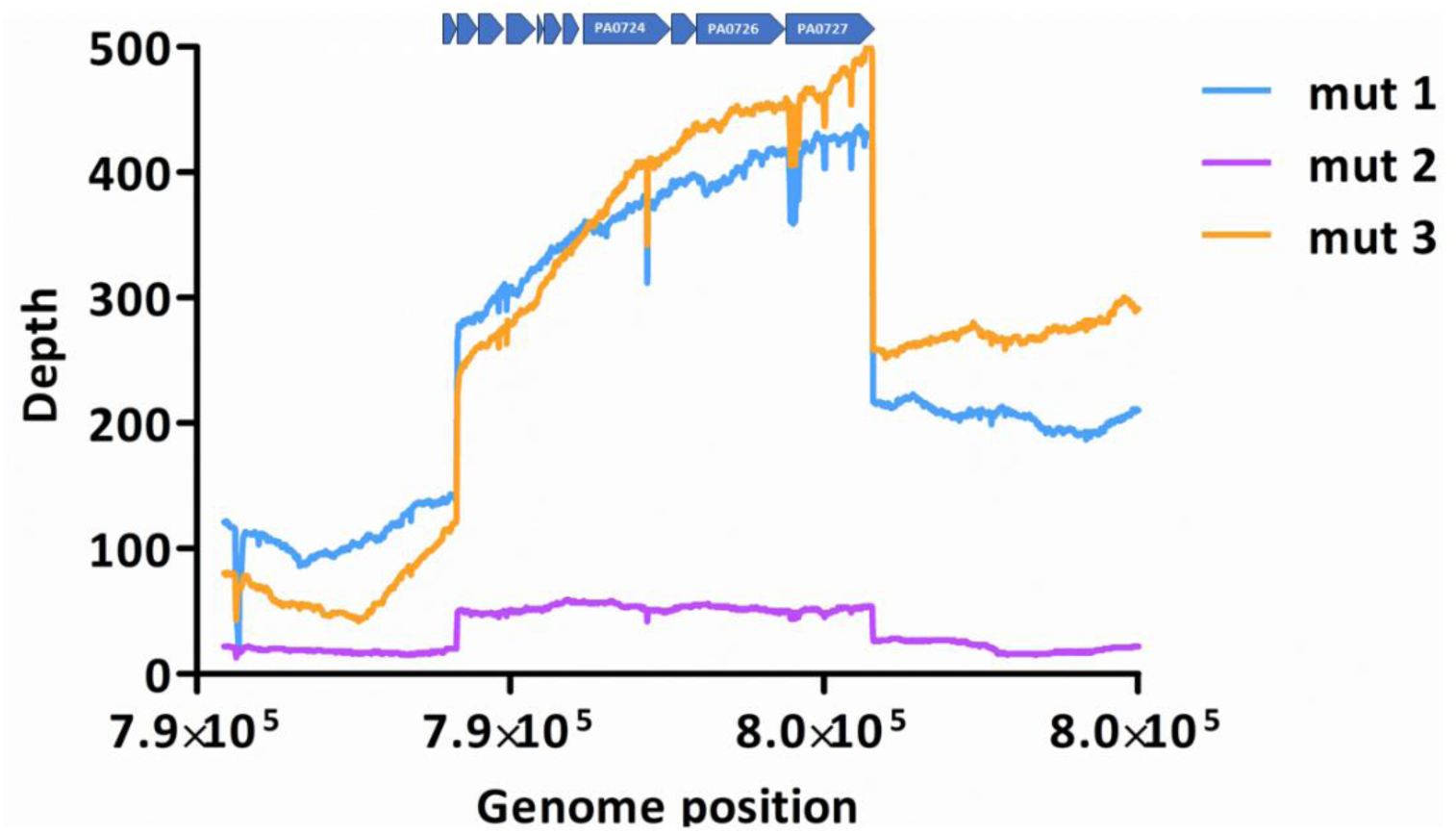
Overexpression of *srkA* null activates prophage Pf4 in *P. aeruginosa*. The depth graph shows the distribution of sequencing read coverage across the genomic region spanning the Pf4 cluster (PA0717-PA0727), with each gene represented by a blue arrow. Three PAO1 WT-derived mutants overexpressing *srkA* null are shown.

**S5 Fig.**
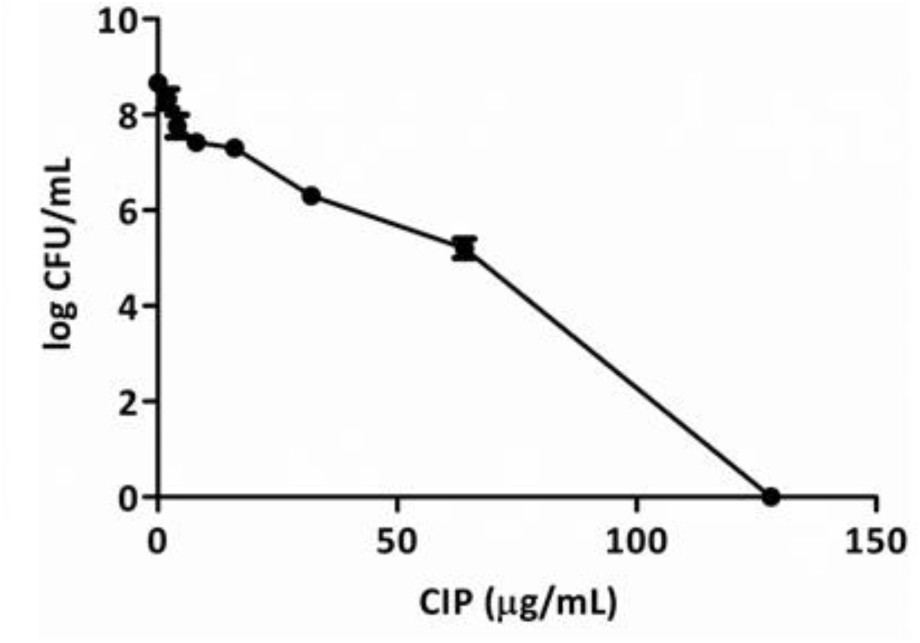
**MIC of PAO1 wild type in the EVPL Model**. Log-transformed total CFU of *P. aeruginosa* PAO1 WT recovered from the EVPL model following treatment with different concentrations of CIP. Three biological replicates were grown on EVPL tissue for 48 h, then exposed to CIP or PBS as a control for 18 h. CFU/lung was determined post treatment. The MIC of CIP in this model was 128 µg/mL.

**S6 Fig.**
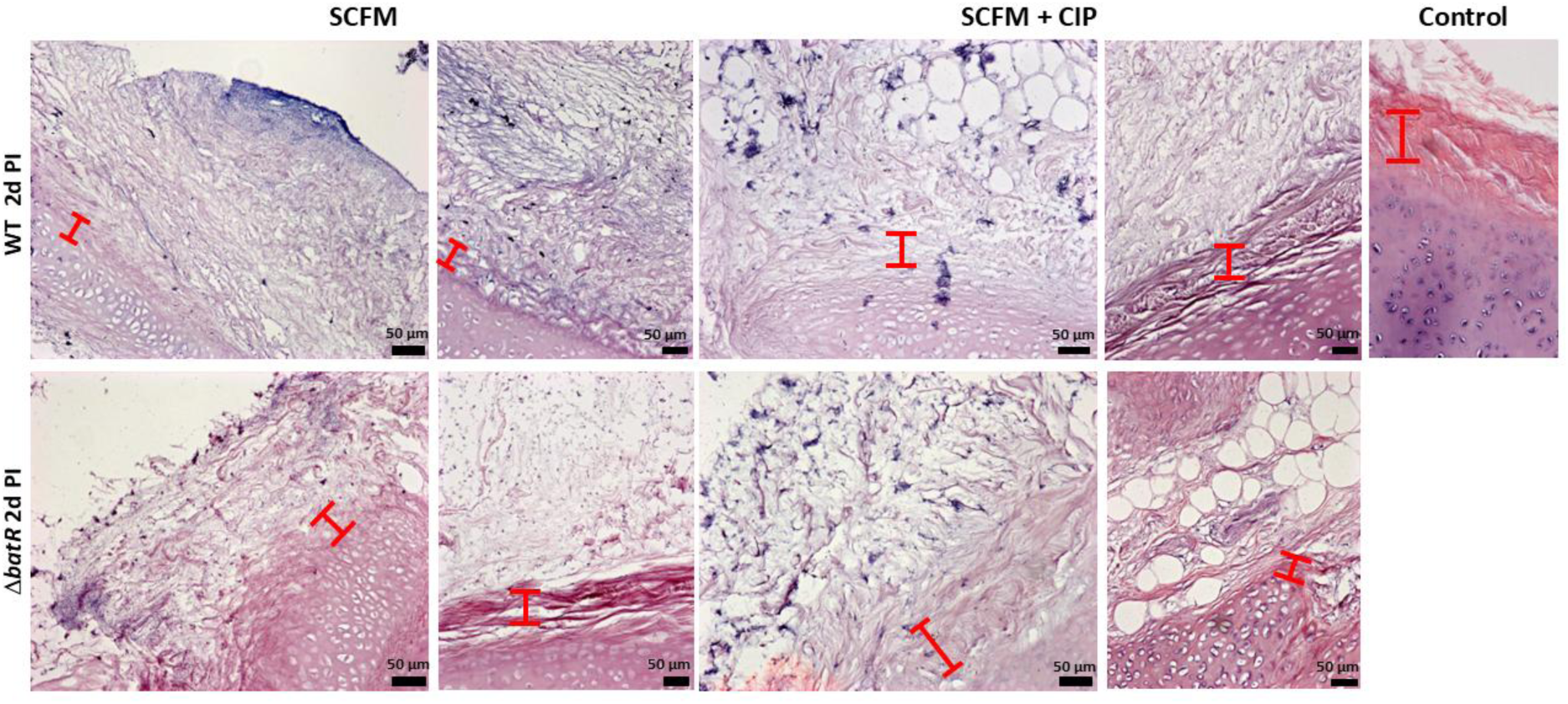
Haematoxylin and eosin (H & E) stained sections of EVPL bronchiolar tissue with SCFM medium infected with *P. aeruginosa* at 2 d post infection. EVPL was infected with *P. aeruginosa* PAO1 WT and Δ*batR*, with uninfected tissue as a negative control. The x20 magnification images from the sections are shown here for non-treated tissues (SCFM) and treated with CIP (SCFM + CIP). Representative images of phenotypes at 2dpi are shown here, but the same results were observed for all biological replicates analysed.

## References

1. Vacheron J, Heiman CM, Keel C. Live cell dynamics of production, explosive release and killing activity of phage tail-like weapons for kin exclusion. Commun Biol. 2021;4(1).

2. Rice LB. Federal funding for the study of antimicrobial resistance in nosocomial pathogens: no ESKAPE. J Infect Dis. 2008;197(8):1079–81.

3. Shigemura K, Arakawa S, Sakai Y, Kinoshita S, Tanaka K, Fujisawa M. Complicated urinary tract infection caused by Pseudomonas aeruginosa in a single institution (1999-2003). Int J Urol. 2006;13(5):538–42.

4. Chitkara YK, Feierabend TC. Endogenous and exogenous infection with Pseudomonas aeruginosa in a burns unit. Int Surg. 1981;66(3):237–40.

5. Mulcahy LR, Isabella VM, Lewis K. Pseudomonas aeruginosa biofilms in disease. Microb Ecol. 2014;68(1):1–12.

6. Kielhofner M, Atmar RL, Hamill RJ, Musher DM. Life-threatening Pseudomonas aeruginosa infections in patients with human immunodeficiency virus infection. Clin Infect Dis. 1992;14(2):403–11.

7. Lyczak JB CC, Pier GB. Lung infections associated with cystic fibrosis. . Clin Microbiol Rev. 2002.

8. Pressler T, Bohmova C, Conway S, Dumcius S, Hjelte L, Høiby N, et al. Chronic *Pseudomonas aeruginosa* infection definition: EuroCareCF Working Group report. Journal of Cystic Fibrosis. 2011;10:S75–S8.

9. Boyd CD, O’Toole GA. Second messenger regulation of biofilm formation: breakthroughs in understanding c-di-GMP effector systems. Annu Rev Cell Dev Biol. 2012;28:439–62.

10. Serra DO, Hengge R. Stress responses go three dimensional - the spatial order of physiological differentiation in bacterial macrocolony biofilms. Environ Microbiol. 2014;16(6):1455–71.

11. Piazza A, Parra L, Ciancio Casalini L, Sisti F, Fernandez J, Malone JG, et al. Cyclic di-GMP Signaling Links Biofilm Formation and Mn(II) Oxidation in Pseudomonas resinovorans. mBio. 2022;13(6):e0273422.

12. Williamson KS, Richards LA, Perez-Osorio AC, Pitts B, McInnerney K, Stewart PS, et al. Heterogeneity in Pseudomonas aeruginosa biofilms includes expression of ribosome hibernation factors in the antibiotic-tolerant subpopulation and hypoxia-induced stress response in the metabolically active population. J Bacteriol. 2012;194(8):2062–73.

13. Fajardo A, Martínez JL. Antibiotics as signals that trigger specific bacterial responses. Current Opinion in Microbiology. 2008;11(2):161–7.

14. Davies J, Spiegelman GB, Yim G. The world of subinhibitory antibiotic concentrations. Curr Opin Microbiol. 2006;9(5):445–53.

15. Ranieri MR, Whitchurch CB, Burrows LL. Mechanisms of biofilm stimulation by subinhibitory concentrations of antimicrobials. Curr Opin Microbiol. 2018;45:164–9.

16. Wright EA, Fothergill JL, Paterson S, Brockhurst MA, Winstanley C. Sub-inhibitory concentrations of some antibiotics can drive diversification of populations in artificial sputum medium. Bmc Microbiol. 2013;13.

17. Taccetti G, Francalanci M, Pizzamiglio G, Messore B, Carnovale V, Cimino G, et al. Cystic Fibrosis: Recent Insights into Inhaled Antibiotic Treatment and Future Perspectives. Antibiotics-Basel. 2021;10(3).

18. Dwyer DJ, Belenky PA, Yang JH, MacDonald IC, Martell JD, Takahashi N, et al. Antibiotics induce redox-related physiological alterations as part of their lethality. P Natl Acad Sci USA. 2014;111(20):E2100–E9.

19. Linares JF, Gustafsson I, Baquero F, Martinez JL. Antibiotics as intermicrobial signaling agents instead of weapons. Proc Natl Acad Sci U S A. 2006;103(51):19484–9.

20. Nalca Y, Jansch L, Bredenbruch F, Geffers R, Buer J, Haussler S. Quorum-sensing antagonistic activities of azithromycin in Pseudomonas aeruginosa PAO1: a global approach. Antimicrob Agents Chemother. 2006;50(5):1680–8.

21. Hoffman LR, D’Argenio DA, MacCoss MJ, Zhang Z, Jones RA, Miller SI. Aminoglycoside antibiotics induce bacterial biofilm formation. Nature. 2005;436(7054):1171–5.

22. Kaplan JB. Antibiotic-induced biofilm formation. Int J Artif Organs. 2011;34(9):737–51.

23. Whitchurch CB, Tolker-Nielsen T, Ragas PC, Mattick JS. Extracellular DNA required for bacterial biofilm formation. Science. 2002;295(5559):1487.

24. Turnbull L, Toyofuku M, Hynen AL, Kurosawa M, Pessi G, Petty NK, et al. Explosive cell lysis as a mechanism for the biogenesis of bacterial membrane vesicles and biofilms. Nat Commun. 2016;7:11220.

25. Petrova OE, Schurr JR, Schurr MJ, Sauer K. The novel Pseudomonas aeruginosa two-component regulator BfmR controls bacteriophage-mediated lysis and DNA release during biofilm development through PhdA. Mol Microbiol. 2011;81(3):767–83.

26. Allesen-Holm M, Barken KB, Yang L, Klausen M, Webb JS, Kjelleberg S, et al. A characterization of DNA release in Pseudomonas aeruginosa cultures and biofilms. Mol Microbiol. 2006;59(4):1114–28.

27. Das T, Manefield M. Pyocyanin Promotes Extracellular DNA Release in. Plos One. 2012;7(10).

28. Wilson R, Sykes DA, Watson D, Rutman A, Taylor GW, Cole PJ. Measurement of Pseudomonas aeruginosa phenazine pigments in sputum and assessment of their contribution to sputum sol toxicity for respiratory epithelium. Infect Immun. 1988;56(9):2515–7.

29. Kaleta MF, Petrova OE, Zampaloni C, Garcia-Alcalde F, Parker M, Sauer K. A previously uncharacterized gene, PA2146, contributes to biofilm formation and drug tolerance across the ɣ- Proteobacteria. NPJ Biofilms Microbiomes. 2022;8(1):54.

30. Yamagishi M, Matsushima H, Wada A, Sakagami M, Fujita N, Ishihama A. Regulation of the Escherichia coli rmf gene encoding the ribosome modulation factor: growth phase- and growth rate-dependent control. EMBO J. 1993;12(2):625–30.

31. Wada A, Yamazaki Y, Fujita N, Ishihama A. Structure and probable genetic location of a "ribosome modulation factor" associated with 100S ribosomes in stationary-phase Escherichia coli cells. Proc Natl Acad Sci U S A. 1990;87(7):2657–61.

32. Akiyama T, Williamson KS, Schaefer R, Pratt S, Chang CB, Franklin MJ. Resuscitation of Pseudomonas aeruginosa from dormancy requires hibernation promoting factor (PA4463) for ribosome preservation. Proc Natl Acad Sci U S A. 2017;114(12):3204–9.

33. Harrison F, Muruli A, Higgins S, Diggle SP. Development of an ex vivo porcine lung model for studying growth, virulence, and signaling of Pseudomonas aeruginosa. Infect Immun. 2014;82(8):3312–23.

34. Werthen M, Henriksson L, Jensen PO, Sternberg C, Givskov M, Bjarnsholt T. An in vitro model of bacterial infections in wounds and other soft tissues. APMIS. 2010;118(2):156–64.

35. Abramson J, Adler J, Dunger J, Evans R, Green T, Pritzel A, et al. Accurate structure prediction of biomolecular interactions with AlphaFold 3. Nature. 2024;630(8016):493–500.

36. Prossliner T, Skovbo Winther K, Sorensen MA, Gerdes K. Ribosome Hibernation. Annu Rev Genet. 2018;52:321–48.

37. Susc ECA. Determination of minimum inhibitory concentrations (MICs) of antibacterial agents by agar dilution. Clin Microbiol Infec. 2000;6(9):509–15.

38. Trampari E, Holden ER, Wickham GJ, Ravi A, Martins LO, Savva GM, et al. Exposure of Salmonella biofilms to antibiotic concentrations rapidly selects resistance with collateral tradeoffs. NPJ Biofilms Microbiomes. 2021;7(1):3.

39. Marti M, Frigols B, Serrano-Aroca A. Antimicrobial Characterization of Advanced Materials for Bioengineering Applications. J Vis Exp. 2018(138).

40. Reales-Calderon JA, Sun Z, Mascaraque V, Perez-Navarro E, Vialas V, Deutsch EW, et al. A wide-ranging Pseudomonas aeruginosa PeptideAtlas build: A useful proteomic resource for a versatile pathogen. J Proteomics. 2021;239:104192.

41. Nakayama K, Takashima K, Ishihara H, Shinomiya T, Kageyama M, Kanaya S, et al. The R-type pyocin of Pseudomonas aeruginosa is related to P2 phage, and the F-type is related to lambda phage. Mol Microbiol. 2000;38(2):213–31.

42. Toyofuku M, Nomura N, Eberl L. Types and origins of bacterial membrane vesicles. Nat Rev Microbiol. 2019;17(1):13–24.

43. Schlimpert S, Flardh K, Buttner J. Fluorescence Time-lapse Imaging of the Complete S. venezuelae Life Cycle Using a Microfluidic Device. J Vis Exp. 2016(108):53863.

44. Dorsey-Oresto A, Lu T, Mosel M, Wang X, Salz T, Drlica K, et al. YihE kinase is a central regulator of programmed cell death in bacteria. Cell Rep. 2013;3(2):528–37.

45. Karimova G, Pidoux J, Ullmann A, Ladant D. A bacterial two-hybrid system based on a reconstituted signal transduction pathway. Proc Natl Acad Sci U S A. 1998;95(10):5752–6.

46. Zheng J, He C, Singh VK, Martin NL, Jia Z. Crystal structure of a novel prokaryotic Ser/Thr kinase and its implication in the Cpx stress response pathway. Mol Microbiol. 2007;63(5):1360–71.

47. Qin S, Xiao W, Zhou C, Pu Q, Deng X, Lan L, et al. Pseudomonas aeruginosa: pathogenesis, virulence factors, antibiotic resistance, interaction with host, technology advances and emerging therapeutics. Signal Transduct Target Ther. 2022;7(1):199.

48. Webb JS, Thompson LS, James S, Charlton T, Tolker-Nielsen T, Koch B, et al. Cell death in Pseudomonas aeruginosa biofilm development. J Bacteriol. 2003;185(15):4585–92.

49. Rice SA, Tan CH, Mikkelsen PJ, Kung V, Woo J, Tay M, et al. The biofilm life cycle and virulence of Pseudomonas aeruginosa are dependent on a filamentous prophage. ISME J. 2009;3(3):271–82.

50. Tortuel D, Tahrioui A, Rodrigues S, Cambronel M, Boukerb AM, Maillot O, et al. Activation of the Cell Wall Stress Response in Pseudomonas aeruginosa Infected by a Pf4 Phage Variant. Microorganisms. 2020;8(11).

51. Harrington NE, Sweeney E, Harrison F. Building a better biofilm - Formation of in vivo-like biofilm structures by Pseudomonas aeruginosa in a porcine model of cystic fibrosis lung infection. Biofilm. 2020;2:100024.

52. Bjarnsholt T JP, Fiandaca MJ, Pedersen J, Hansen CR, Andersen CB, Pressler T, Givskov M, Høiby N. Pseudomonas aeruginosa biofilms in the respiratory tract of cystic fibrosis patients. . Pediatr Pulmonol. 2009.

53. Henderson AG EC, Button B, Abdullah LH, Cai LH, Leigh MW, DeMaria GC, Matsui H, Donaldson SH, Davis CW, Sheehan JK, Boucher RC, Kesimer M. Cystic fibrosis airway secretions exhibit mucin hyperconcentration and increased osmotic pressure. J Clin Invest. 2014.

54. Baltimore RS CC, Smith GJ. Immunohistopathologic localization of Pseudomonas aeruginosa in lungs from patients with cystic fibrosis. Implications for the pathogenesis of progressive lung deterioration. Am Rev Respir Dis. 1989.

55. Bielecki P, Puchalka J, Wos-Oxley ML, Loessner H, Glik J, Kawecki M, et al. In-vivo expression profiling of Pseudomonas aeruginosa infections reveals niche-specific and strain-independent transcriptional programs. PLoS One. 2011;6(9):e24235.

56. Ramos I, Dietrich LE, Price-Whelan A, Newman DK. Phenazines affect biofilm formation by Pseudomonas aeruginosa in similar ways at various scales. Res Microbiol. 2010;161(3):187–91.

57. Price-Whelan A, Dietrich LE, Newman DK. Rethinking ’secondary’ metabolism: physiological roles for phenazine antibiotics. Nat Chem Biol. 2006;2(2):71–8.

58. Das T, Manefield M. Pyocyanin promotes extracellular DNA release in Pseudomonas aeruginosa. PLoS One. 2012;7(10):e46718.

59. Das T, Kutty SK, Kumar N, Manefield M. Pyocyanin facilitates extracellular DNA binding to Pseudomonas aeruginosa influencing cell surface properties and aggregation. PLoS One. 2013;8(3):e58299.

60. Dietrich LE TT, Price-Whelan A, Newman DK. Redox-active antibiotics control gene expression and community behavior in divergent bacteria. Science. 2008.

61. Dietrich LE OC, Price-Whelan A, Sakhtah H, Hunter RC, Newman DK. . Bacterial community morphogenesis is intimately linked to the intracellular redox state. J Bacteriol. 2013.

62. Schiessl KT HF, Jo J, Nazia SZ, Wang B, Price-Whelan A, Min W, Dietrich LEP. Phenazine production promotes antibiotic tolerance and metabolic heterogeneity in Pseudomonas aeruginosa biofilms. Nat Commun. 2019.

63. Garay-Arroyo A, Colmenero-Flores JM, Garciarrubio A, Covarrubias AA. Highly hydrophilic proteins in prokaryotes and eukaryotes are common during conditions of water deficit. J Biol Chem. 2000;275(8):5668–74.

64. Ameisen JC. On the origin, evolution, and nature of programmed cell death: a timeline of four billion years. Cell Death Differ. 2002;9(4):367–93.

65. Rice KC, Bayles KW. Death’s toolbox: examining the molecular components of bacterial programmed cell death. Mol Microbiol. 2003;50(3):729–38.

66. Dietrich LE, Okegbe C, Price-Whelan A, Sakhtah H, Hunter RC, Newman DK. Bacterial community morphogenesis is intimately linked to the intracellular redox state. J Bacteriol. 2013;195(7):1371–80.

67. Miller JH. Experiments in molecular genetics. Cold Spring Harbor, N.Y.: Cold Spring Harbor Laboratory; 1972. xvi, 466 p. p.

68. Holloway BW. Genetic recombination in Pseudomonas aeruginosa. J Gen Microbiol. 1955;13(3):572–81.

69. Woodcock DM, Crowther PJ, Doherty J, Jefferson S, DeCruz E, Noyer-Weidner M, et al. Quantitative evaluation of Escherichia coli host strains for tolerance to cytosine methylation in plasmid and phage recombinants. Nucleic Acids Res. 1989;17(9):3469–78.

70. Scott TA, Heine D, Qin Z, Wilkinson B. An L-threonine transaldolase is required for L-threo-beta-hydroxy-alpha-amino acid assembly during obafluorin biosynthesis. Nat Commun. 2017;8:15935.

71. Heeb S, Itoh Y, Nishijyo T, Schnider U, Keel C, Wade J, et al. Small, stable shuttle vectors based on the minimal pVS1 replicon for use in gram-negative, plant-associated bacteria. Mol Plant Microbe Interact. 2000;13(2):232–7.

72. Choi KH, Gaynor JB, White KG, Lopez C, Bosio CM, Karkhoff-Schweizer RR, et al. A Tn7-based broad-range bacterial cloning and expression system. Nat Methods. 2005;2(6):443–8.

73. Woodcock SD, Syson K, Little RH, Ward D, Sifouna D, Brown JKM, et al. Trehalose and alpha-glucan mediate distinct abiotic stress responses in Pseudomonas aeruginosa. PLoS Genet. 2021;17(4):e1009524.

74. Piazza A, Ciancio Casalini L, Pacini VA, Sanguinetti G, Ottado J, Gottig N. Environmental Bacteria Involved in Manganese(II) Oxidation and Removal From Groundwater. Front Microbiol. 2019;10:119.

75. Trampari E, Zhang C, Gotts K, Savva GM, Bavro VN, Webber M. Cefotaxime Exposure Selects Mutations within the CA-Domain of envZ Which Promote Antibiotic Resistance but Repress Biofilm Formation in Salmonella. Microbiol Spectr. 2022;10(3):e0214521.

76. Thompson CMA, Hall JPJ, Chandra G, Martins C, Saalbach G, Panturat S, et al. Plasmids manipulate bacterial behaviour through translational regulatory crosstalk. PLoS Biol. 2023;21(2):e3001988.

77. Schindelin J, Arganda-Carreras I, Frise E, Kaynig V, Longair M, Pietzsch T, et al. Fiji: an open-source platform for biological-image analysis. Nat Methods. 2012;9(7):676–82.

78. Cornell WC, Morgan CJ, Koyama L, Sakhtah H, Mansfield JH, Dietrich LEP. Paraffin Embedding and Thin Sectioning of Microbial Colony Biofilms for Microscopic Analysis. J Vis Exp. 2018(133).

79. Deatherage DE, Barrick JE. Identification of mutations in laboratory-evolved microbes from next-generation sequencing data using breseq. Methods Mol Biol. 2014;1151:165–88.

80. Robinson JT, Thorvaldsdottir H, Winckler W, Guttman M, Lander ES, Getz G, et al. Integrative genomics viewer. Nat Biotechnol. 2011;29(1):24–6.

81. Harrington NE, Sweeney E, Alav I, Allen F, Moat J, Harrison F. Antibiotic Efficacy Testing in an Ex vivo Model of Pseudomonas aeruginosa and Staphylococcus aureus Biofilms in the Cystic Fibrosis Lung. J Vis Exp. 2021(167).

82. Perez-Riverol Y, Bai J, Bandla C, Garcia-Seisdedos D, Hewapathirana S, Kamatchinathan S, et al. The PRIDE database resources in 2022: a hub for mass spectrometry-based proteomics evidences. Nucleic Acids Res. 2022;50(D1):D543–D52.

